# Phospholipase D3 regulates lysosomal morphology, biogenesis, and function in neurons

**DOI:** 10.1101/2024.09.26.615214

**Authors:** Yongchao Wang, Cristian Carvajal-Tapia, Stuti Jain, Emmeline Wang, Paige Rieschick, Nathaniel L. Hepowit, Nathan Appelbaum, Alex Prusky, Theodora Anghelina, Avelyn Kulsomphob, Vaibhav A. Janve, Samuel J. Frank, Alena Shostak, Timothy J. Hohman, Bret C. Mobley, Jason A. MacGurn, Wilber Romero-Fernandez, Matthew S. Schrag

**Affiliations:** Department of Neurology, Vanderbilt University Medical Center, Nashville, Tennessee 37240, USA; Department of Cell and Developmental Biology, Vanderbilt University, Nashville, Tennessee 37240, USA; Vanderbilt Genetics Institute, Vanderbilt University Medical Center, Nashville, Tennessee 37232, USA; Vanderbilt Memory and Alzheimer’s Center, Vanderbilt University, Nashville, Tennessee 37235, USA; Department of Pathology, Microbiology, and Immunology, Vanderbilt University Medical Center, Nashville, Tennessee, USA

**Keywords:** Alzheimer’s Disease, Phospholipase D3, Transcription factor EB, transcription factor binding to IGHM enhancer 3, Cathepsin B, Ubiquitin, Ubiquitination, Lysosome biogenesis

## Abstract

Phospholipase D3 (PLD3) is a lysosomal protein in neurons and is linked to an Alzheimer’s disease (AD) risk through coding variants. Its role in lysosomal abnormalities observed in AD remains unclear. We conducted a comprehensive study integrating transcriptomic, proteomic, and cell biology approaches to investigate PLD3’s role in lysosome function in neurons.

PLD3 undergoes aberrant ubiquitination in AD brain tissue, leading to altered protein abundance and subcellular location. We assessed the lysosomal proteome in SH-SY5Y cells with PLD3 knockout and found PLD3^−/−^ cells had higher levels of proteins related to lysosomal biogenesis, endocytosis, and calcium signaling. Loss of PLD3 caused an enlargement in neuronal lysosome size, increased endocytic activity and a decline in lysosomal clearance of aggregated human β-amyloid or α-synuclein, despite increased expression level and activity of cathepsin B. PLD3-deficiency increase the level and transcriptional activity of transcription factor EB (TFEB) and transcription factor binding to IGHM enhancer 3 (TFE3), suggesting a role for PLD3 in regulating lysosomal biogenesis.

In summary, we found that PLD3 is required to maintain neuronal lysosomal function.

## Background

Alzheimer’s disease (AD) is the most common cause of dementia. It is characterized by a broad range of neuropathologies, including extracellular accumulation of β-amyloid (Aβ) in plaques and cerebral vessels, intraneuronal tangles and neuropil threads composed of hyperphosphorylated tau, dystrophic neurites, granulovacuolar degenerating bodies, and synaptic loss (1, 2).

Degradation of waste proteins in neurons is managed by the concerted action of the ubiquitin-proteasome (UPS) (3) and the endosome-autophagosome-lysosome pathways (EAL) (4). If these clearance pathways are dysregulated, proteins may accumulate and impair cellular homeostasis. Aberrant ubiquitination and dysfunction in UPS have long been associated with AD (5, 6).

Many studies have implicated sub-optimal function of lysosomal machinery in the pathogenesis of neurodegenerative proteinopathies (7, 8). Various factors, including alterations in lysosome size, population or trafficking, pH regulation, EAL disruption, and lysosomal membrane permeabilization, can cause lysosomes to malfunction (9, 10). Lysosomal deacidification, which could contribute to inefficient function of the more than 60 lysosomal hydrolases, has been hypothesized to play a role in the pathogenesis of several neurodegenerative diseases (11, 12).

Lysosomal biogenesis is regulated by Transcription factor EB (TFEB) and transcription factor binding to IGHM enhancer 3 (TFE3) levels and activity, because TFEB and TFE3 govern the expression of a network of genes linked to lysosomal biogenesis and autophagy (13–16). Alterations in the TFEB/TFE3 pathway have been observed in AD *(for a review, see* (*15*)). It has been reported that genetic disruption of TFEB results in defective microglia activation and increased tau pathology (17).

Restoring lysosomal homeostasis would likely be beneficial for treating AD, so understanding the role of lysosomal proteins like PLD3, which is linked to AD risk, is a key priority.

PLD3 is a member of the phospholipase D family and is primarily localized to lysosomes. It is highly expressed in neurons (18, 19). A growing body of evidence has linked PLD3 to AD (20, 21). PLD3 is remarkably enriched in dystrophic neurites surrounding ß-amyloid plaques in AD(21).

Axonal PLD3 accumulation was functionally implicated in the formation and growth of dystrophic neurites, contributing to neural network defects in AD (22). Lysosomes in neurons of PLD3 knockout mice exhibited significant enlargement (23). Despite its localization in neuronal lysosomes, its role(s) in lysosomal function remains a matter of ongoing study, with recent evidence pointing to a role in lysosomal degradation of lipids and nucleic acids (24, 25).

Our study presents evidence supporting a causal link between PLD3 and lysosome abnormalities occurring in AD, including morphological changes and impairment of degradative function that can promote proteostasis disturbance. Additionally, we demonstrate that PLD3 regulates TFEB/TFE3 signaling, suggesting a role in lysosomal biogenesis.

## Methods

### Human brain tissue

Brain tissue was obtained from the Vanderbilt Brain and Biospecimen Bank at Vanderbilt University Medical Center, Nashville, Tennessee, USA (IRB# 180287). Written informed consent for each anatomical donation was obtained from patients or their surrogate decision-makers. The Vanderbilt University Medical Center Institutional Review Board provided ethical supervision, and the study was performed in accordance with the ethical principles of the Declaration of Helsinki.

### Antibodies and reagents

Primary antibodies used in this study include: rabbit anti-PLD3 (HPA012800, Sigma-Aldrich, St. Louis, MO, 1:1000 for WB and 10 μg/mL for IHC), mouse anti-ubiquitin clone Ubi-1 (13-1600, Invitrogen, Carlsbad, CA, 1:1000 for WB and 10 μg/mL for IHC), mouse anti-TFEB (91767, Cell Signaling Technology, Boston, MA, 1:1000 for WB, 1:100 for IF), rabbit anti-TFE3 (81744, Cell Signaling Technology, 1:1000 for WB), mouse anti-Lamin B1 (66095, Proteintech, Rosemont, IL, 1:1000 for WB), mouse anti-LC3B (83506, Cell Signaling Technology, 1:1000 for WB), mouse anti-LAMP2 (H4B4, DSHB, Iowa, IA, 1:1000 for WB and 10 μg/mL for IHC), mouse anti-GAPDH (ab59164, Abcam, Cambridge, UK, 1:5000 for WB), mouse anti-GAPDH (60004-1-Ig; Proteintech, 1:1000 for WB), mouse anti-GAPDH-HRP (60004; Proteintech, 1:1000 for WB), rabbit anti-β-Tubulin (15115, Cell Signaling Technology, 1:5000 for WB), goat anti-CTSB (AF965, R&D Systems, Minneapolis, MN, 1:1000 for WB and 10 μg/mL for IHC), goat anti-CTSD (AF1014, R&D systems, 1:1000 for WB and 10 μg/mL for IHC), rabbit anti-IKKβ (8943, Cell Signaling Technology, 1:1000 for WB), normal rabbit IgG (12-370, Cell Signaling Technology).

Secondary antibodies used in this study include: goat anti-mouse IgG (H+L), HRP (62-6520; Invitrogen), goat anti-rabbit IgG (H+L), HRP (65-6120; Invitrogen), IRDye® 800 CW goat anti-rabbit (926-32211; LI-COR, Lincoln, NE), IRDye® 680 RD goat anti-mouse (926-68070; LI-COR), IRDye® 680 LT donkey anti-goat (926-68024; LI-COR), IRDye® 800 CW donkey anti-rabbit (926-32213; LI-COR), and IRDye® 800 CW donkey anti-mouse (926-32213; LI-COR).

Reagents used in this study include: synthetic human β-amyloid_1-42_ (AG968; Sigma-Aldrich), DQ Green BSA (D12050, ThermoFisher Scientific), dextran Oregon Green 488 (D7172, ThermoFisher Scientific), Protein G/A agarose supernatant (IP05, Millipore, Burlington, MA), dextran Alexa Fluor 594 (D22913, ThermoFisher Scientific), dextran Alexa Fluor 647 (D22914, ThermoFisher Scientific), Alexa Fluor 647 BSA (A34785, ThermoFisher Scientific), CellLight Lysosome-RFP BacMam 2.0 (C10597, ThermoFisher Scientific), Torin 1 (14379, Cell Signaling Technology), Hoechst 33342 (R37605, ThermoFisher Scientific, two drops/ml), neurobasal-A medium, minus phenol red (12349015, ThermoFisher Scientific), 1 µM 4,4’-[(2-methoxy-1,4-phenylene) di-(1*E*)-2,1-ethenediyl] bisphenol (MX-04) (Tocris, Minneapolis, MN), Thiazine Red (1 µM, Chemsavers Inc, Bluefield, VA), 4’, 6-diamidino-2-phenylindol (1 µM, DAPI, Cayman Chemical, Ann Arbor, MI), TO-PRO™-3 (1 µM, ThermoFisher Scientific). Potent and specific CTSB inhibitor CA074 (1 µM, 4863, Tocris). The information for all other reagents is indicated in the corresponding sections. Details of specific methods are provided in the supplementary material.

### Brain tissue preparation, immunostaining, and proximity ligation assay

Human brain tissue was prepared as previously described (26). Immunohistochemical labeling was performed as described (21, 26–28). PLD3-ubiquitin complex detection in postmortem human brain was performed using the Duolink® Proximity Ligation Assay (PLA) kit (Sigma-Aldrich), following the protocol previously described (29, 30), including modifications optimized for a fluorescent format in the human brain (26).

### Human Aβ seeds extraction

Human Aβ seeds were isolated as previously described (31) with minor modifications. Seeds were isolated from the brain tissue of patients with severe cerebral amyloid angiopathy (CAA) with minimal other neuropathology. Briefly, tissue was homogenized, and the lipid-rich fraction was removed by centrifuging in 17% dextran (40,000 MW, Sigma-Aldrich) at 10,000 x g for 5 minutes. The pellet was resuspended in TBS (Corning, Corning, NY), and the suspension was filtered through a custom-designed 3D printed microsieve apparatus with 40 µM pore size to isolate microvascular fragments. Aggregates of β-amyloid were freed from the vascular matrix by incubating with 3 mg/ml Collagenase Type I (Gibco, Billings, MT) overnight at 37°C and gently homogenizing with 12 passes of a Dounce homogenizer. The suspension was again passed through the microsieve, and the liquid containing the particles was centrifuged and resuspended in TBS (Corning). It was pelleted and resuspended twice more to remove any residual enzyme. The particles were disrupted with sonication to produce a microparticulate suspension with a mean particle diameter of around 10 microns (particle size distribution shown in Fig. S7). Before adding to cells, particles were further disrupted with sonication to produce a microparticulate suspension with a mean particle diameter of about 1 micron (Fig. S7).

### PLD3 knockout cell line

SH-SY5Y cell line (CRL-2266, ATCC) was used to generate PLD3 knockout (SH-SY5Y PLD3^−/−^), performed by Abcam (Abcam, Waltham, MA). PLD3-g1: AGCCCACAACCGCCAGAATG and PLD3-g2: TACTCGCAAGGGTCATAGCA were transfected into SH-SY5Y cells individually with the Cas9 gene to generate the PLD3 knockout two PLD3 knockout clone 3 (PLD3*^−/−^*) and clone 14 PLD3*^−/−c14^*). A DNA deletion (∼100 bp) in exon 5 of the PLD3 gene in both alleles was achieved, making the frameshift knockout. PCR subsequently confirmed the resulting DNA deletion and was further validated at the protein level by western blotting. Unless otherwise specified, all experiments were performed with clone 3 (PLD3^−/−^).

### Cell culture, cDNA transfection, siRNA, differentiation to neurons, and drug incubation

SH-SY5Y wild type (WT, CRL 2266, ATCC) and SH-SY5Y PLD3^−/−^ cells were grown in DMEM/F12 (Gibco) supplemented with 10% (v/v) fetal bovine serum (FBS, Corning) and 100 μg/mL Normocin (InvivoGen) or 1% penicillin/streptomycin (0503, ScienceCell, Carlsbad, CA) at 37°C in a humidified atmosphere of 5% CO_2_.

PLD3 plasmids were from Sino Biological (Beijing, China), and the PLD3 mutant plasmids were previously generated using a site-directed mutagenesis kit (21). GFP-N-TFEB (38119, Addgene) and its corresponding control (60360, Addgene) plasmids were from Addgene. PLD3 siRNAs and their corresponding control siRNAs were purchased from IDT. For plasmid transfection, plasmid DNA was transfected into cells with Lipofectamine 3000 reagent following the manufacturer’s protocol (L3000015, Invitrogen). For siRNA transfection, cells were transfected with 10 μM siRNA by Oligofectamine^TM^ reagent per the manufacturer’s recommendations (12252-011, Invitrogen). SH-SY5Y differentiation into neurons was performed as previously described (32).

Cells were incubated with and without synthetic β-amyloid_1-42_ (1 µg/mL) for 1, 3, 5, or 7 days before treatment with DMSO, dextran (0.25 mg/mL), HBSS, or Torin 1 (1 µM) for 3 hours at 37℃, followed by washing with PBS for 10 minutes. The cells were then fixed following immunocytochemistry as described above. Subcellular fluorescence was detected, imaged, and analyzed using the CellInsight CX7 LED Pro HCS Platform.

### Human induced pluripotent stem cell (iPSC) maintenance and differentiation to neurons

CC3 iPSCs (33) were maintained and differentiated into cortical glutamatergic neurons as previously described (34) with minor modifications (35). iPSC neurons in culture were incubated with and without human β-amyloid seeds (100 ng/mL) for 2, 3, 5, or 7 days before performing protein extraction, SDS-PAGE, and western blot as described below, or immunocytochemistry as described above.

### Cathepsin B activity measurement

Cells were plated at a concentration of 5 × 10^^6^ cells/75 cm^2^ flasks and cultured overnight before assay. Cells were collected, and the CTSB enzymatic activity was measured according to the guidelines of SensoLyte® 520 Cathepsin B Assay Kit Fluorometric (AS-72164; AnaSpec, Fremont, CA). Results were normalized to the total protein concentration determined by Pierce™ BCA Protein Assay Kit (23225; ThermoFisher Scientific) and enzymatic activity in the presence of the potent and selective inhibitor of CTSB, CA074 (1 μM).

### Protein extraction, SDS-PAGE, and western blot

Cell protein homogenates, and grey matter, and white matter tissue lysates were prepared as described previously (21). SDS-PAGE and western blot were performed as described previously (21).

### Lysosome labeling with dextran

Cells were seeded at a concentration of 40 × 10^^3^ cells per well in Nunc microwell 96-well optical-bottom plates and allowed to attach overnight. The cells were then incubated with dextran Oregon Green (0.25 mg/mL) and dextran AlexaFluor 594 (0.25 mg/mL) overnight at 37℃, followed by a chasing period of 6 hours to minimize endosome labeling. The staining medium was removed, and the cells were incubated with Neurobasal medium containing Hoechst for 2 hours at 37℃. The subcellular fluorescence patterns were quantified using CellInsight CX7 and the LED Pro HCS Platform.

### Co-localization analysis of dextran and LAMP1-RFP

Cells were seeded at 40 × 10^^3^ cells per well in Nunc microwell 96-well optical-bottom plates and allowed to attach overnight. A construct expressing LAMP1-fused RFP was delivered into cells using BacMam 2.0 technology to target lysosomes. On the second day, the medium was removed, and the cells were incubated with dextran AlexaFluor 647 (0.25 mg/mL) overnight in the incubator, followed by a chasing period of 6 hours. The staining medium was removed, and the cells were stained with Hoechst in neurobasal-A medium with clear color for 2 hours at 37℃. The subcellular fluorescence patterns were quantified using CellInsight CX7 and the LED Pro HCS Platform.

### Lysosomal function assay

Cells were seeded at 40 × 10^^3^ cells per well in Nunc microwell 96-well optical-bottom plates and allowed to attach overnight. The cells were then treated with DMSO, Torin-1 (1 μM), or Bafilomycin A (100 nM) for 3 hours, followed by incubation with DQ Green BSA (10 μg/mL) and Alexa Fluor 647 BSA (10 μg/mL) for 3 hours. The assay medium was removed, and the cells were incubated with Hoechst in neurobasal-A medium for nuclear staining for 2 hours at 37°C. Subcellular fluorescence was subsequently analyzed using the CellInsight CX7 LED Pro HCS Platform.

### Spot detection and quantification by CellInsight CX7 LED Pro HCS platform

Subcellular fluorescence was detected and imaged in acquisition mode on the CX7 HCS platform. Images *(n=30-50)* per well were automatically captured using laser autofocus, and the in-focus pictures were selected for subsequent analysis. For quantification analysis, images were first preprocessed with background removal, smoothing, and segmentation, followed by spot (fluorescent signals) identification and validation. The spots were compartmentalized into the cell body part (circle) and the distal part (ring). All images were re-scanned in disk mode, and the information about spot area and intensity (total and average) on all channels was reported and statistically analyzed. Three to six independent experiments *(n=3-6)* were performed for final statistical analysis.

### Transmission electron microscopy

Cells were fixed in 0.1 M cacodylate buffer containing 2.5% glutaraldehyde, harvested using a cell scraper, and pelleted. Samples were then sequentially postfixed in 1% tannic acid and 1% OsO4, followed by en block staining in 2% uranyl acetate. After post-fixation, the samples were dehydrated in a graded ethanol series and infiltrated with Quetol 651-based Spurr’s resin using propylene oxide as a transition solvent. The resin was polymerized at 60 °C for 48 hours, and 70 nm sections were cut on a Leica UC7 ultramicrotome.

Imaging was performed on a JEOL 2100+ transmission electron microscope operating at 200 keV, with images captured using an AMT nanosprint CMOS camera. Single images were acquired with AMT imaging software, while tiled image sets were acquired using SerialEM automation software. Tiled datasets were processed using the IMOD/etomo software suite and FIJI. Lysosome characteristics were analyzed and quantified using FIJI.

### Lysosomal protein preparation

Lysosomes were isolated and purified as described previously (21).

### Isolation of cytoplasmic and nuclear proteins

Cytoplasmic *(c)* and nuclear *(n)* proteins were isolated from cells of interest following the instructions in the manufacturer’s user guide (78833, ThermoFisher Scientific).

### Mass spectrometry and bioinformatic analysis

Protein samples were prepared with 5% SDS, processed using S-Trap (ProtiFi) digestion, and analyzed by LC-MS/MS, following a protocol described previously (36). The LC-MS/MS data were analyzed using MaxQuant against a human database created from the UniprotKB protein database, with MaxQuant contaminants added (37). Standard MaxQuant parameters were applied, along with LFQ and match-between-runs. Variable modifications included methionine oxidation and N-terminal acetylation, while carbamidomethyl cysteine was a fixed modification. A false discovery rate (FDR) 0.01 was applied to peptide and protein identifications. Label-free quantitative (LFQ) analysis of the identified proteins was performed using the MSstats R package with default parameters (38).

### RNA extraction, sequencing, and bioinformatic analysis

Total RNA was isolated from cells of interest using RNeasy Mini Plus Kits (74134, Qiagen) following the manufacturer’s protocol, with each group having three replicates. RNA quality control was performed using the Agilent Bioanalyzer, and RNA quantity was determined using the RNA Qubit assay (Q10210, ThermoFisher Scientific).

### Library preparation and RNA sequencing

Library preparation was started with ribosomal RNA removal using the NEBNext® rRNA Depletion kit and 500 ng of input total RNA (E6310L, New England Biolabs). Following rRNA depletion, the RNA samples were subjected to mRNA enrichment using oligo(dT) beads, and subsequent fragmentation was done using divalent cations at an elevated temperature. Libraries were generated by cDNA synthesis with random hexamer primers, RNase H digestion, A-tailing, and adapter ligation. The cDNA libraries were assessed for quality with a Bioanalyzer and quantified using the KAPA Library Quantification Kit according to the manufacturer’s recommendations (KK4824, Roche). The libraries were mixed in equal molar ratios, and the resulting library pools underwent cluster generation using the NovaSeq 6000 System according to the manufacturer’s protocol. A 150 bp paired-end sequencing was performed on the NovaSeq 6000 platform, which was set to target 30M reads per sample. Real-Time Analysis Software (RTA) and NovaSeq Control Software (NCS) (1.8.0; Illumina) were used for base calling. MultiQC (v1.7; Illumina) was used for data quality assessments.

### Bioinformatic analysis

The raw sequencing data (FASTQ files) were subjected to quality and adapter trimming and quality control using the wrapper program, Trim Galore. The GRCh38 human transcriptome from GENCODE was used to build the transcriptome index with the Kallisto program (39). After being processed by Trim Galore, the sequences were mapped and quantified into transcript-level abundance using the Kallisto program (39). Transcript-level abundance was then summarized into gene-level abundance, followed by differential gene expression analysis using DESeq2 in R (40). Heatmap representation of gene-level abundance, principal component analysis (PCA), and volcano plot representation of differentially expressed (DE) genes were all performed and graphed with R. Normalized RNA-Seq quantification was then submitted to the GSEA website for gene set enrichment analysis (GSEA) with c2.cp.kegg_-_legacy.v2023.2.Hs.symbols.gmt as the gene set database (41).

### Statistical analyses

The number of samples (*n*) in each group/experimental condition is indicated in the figure legends. All data were graphed and statistically analyzed using GraphPad software (version 10.0.2, GraphPad Software Inc., San Diego, CA) or R packages unless stated otherwise. For gene set enrichment analysis, statistical tests were based on the Kolmogorov-Smirnov test (41). For RNA-Seq differential gene expression analysis, the data were subject to negative binomial generalized linear model fitting and Wald statistics (40). The parametric unpaired *t*-test or the Welch’s unequal variances *t*-test was used to compare two groups. Ordinary one-way ANOVA was used to compare more than two independent groups, followed by Tukey’s, Dunnett multiple comparisons, or Uncorrected Fisher’s LSD. Non-parametric Kruskal-Wallis tests were performed, followed by Dunn’s multiple comparisons test. Two-way ANOVA tests were performed to assess the individual and interaction effects of two variables, followed by the Dunnett test. Statistical tests were specified and noted in the respective figure legends. Unless stated otherwise, a significant level of 0.05 was used to reject the null hypothesis throughout this article.

## Results

### PLD3 is a target of aberrant ubiquitination in human brain with AD resulting in its altered subcellular location

Immunohistochemistry with a validated antibody (Fig. S1) in brain sections without AD (Braak I-II) showed high immunoreactivity for PLD3 in a punctate pattern in all regions analyzed (Fig. 1A and Fig. S2), consistent with the expected lysosomal subcellular distribution. In sections from patients with AD (Braak III-VI), PLD3 was less abundant in healthy-appearing neurons and was enriched in dystrophic neurites surrounding β-amyloid plaques and in neuropil threads and neurofibrillary tangles (Fig. 1B and Fig. S2).

**Figure 1.**
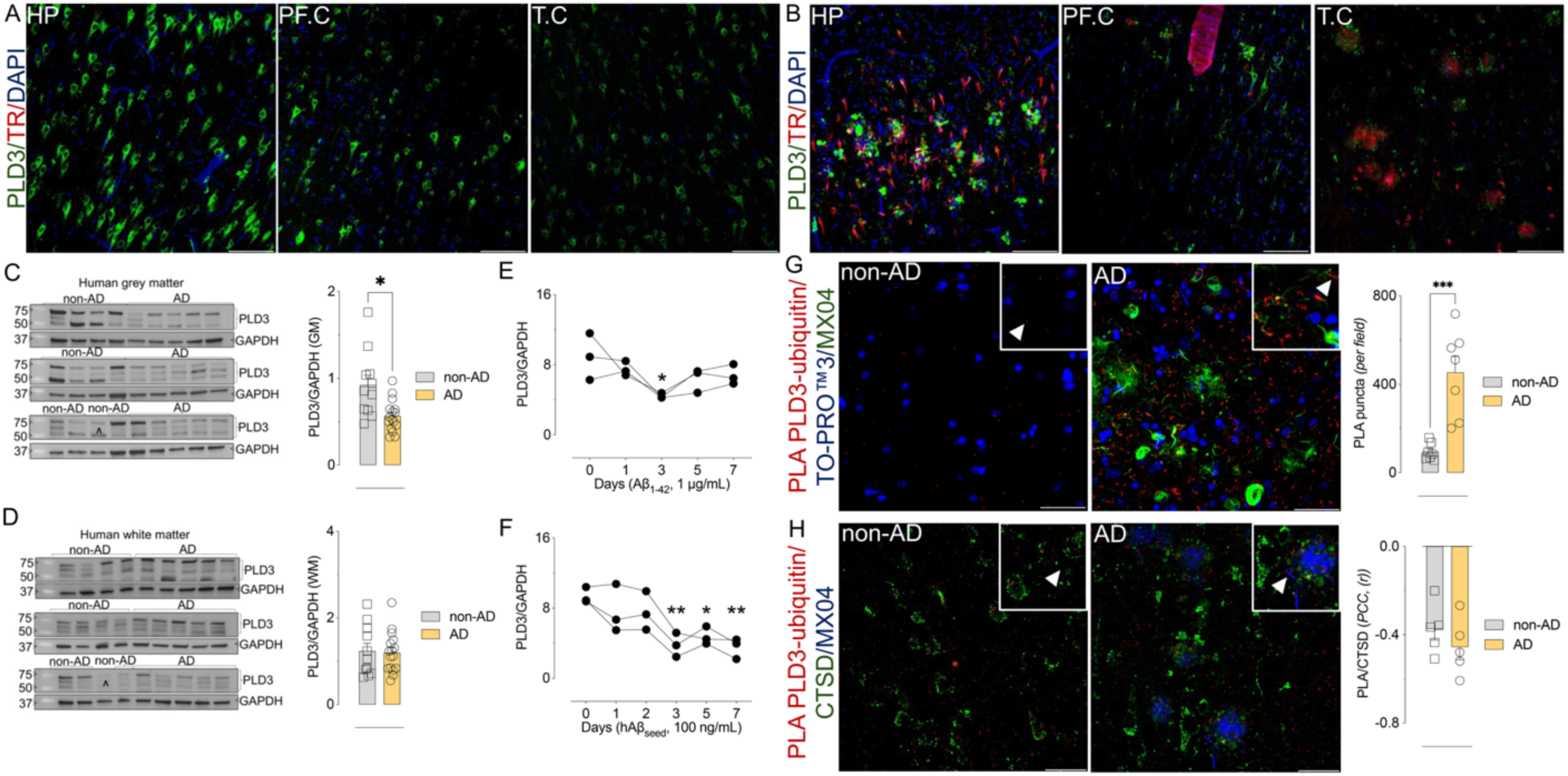
PLD3 is decreased in human AD brain, likely due to ubiquitination. A and B) Immunostaining of PLD3 in the human brain. Representative confocal microscopy images of PLD3 immunostaining *(green)* in the hippocampus (HP), posterior frontal (PF.C), and superior temporal lobe (T.C) sections from (A) non-AD and (B) AD brain sections, *n=8.* Nuclei were stained with DAPI *(blue),* and β-amyloid and neuritic plaques, neurofibrillary tangles, and other tau aggregates were stained with Thiazine Red *(TR, red)*. The scale bars are 100 μm. C and D) PLD3 levels. Western blot analysis of PLD3 in superior temporal lobe lysates from (C) grey matter (GM) and (D) white matter (WM). Data are normalized to GADPH (means ± S.E.M., non-AD *n=11*, and AD *n=16*). \**p* < 0.05 by Welch’s unequal variances *t*-test. E) Outcome of 7-day incubation with synthetic Aβ_1-42_ on PLD3 levels in SY-SY5Y-WT cells. Data presented are normalized to GADPH (mean ± S.E.M., *n=3*). \**p* < 0.05 by one-way ANOVA compared to nontreated cells. α correspond to AD case. F) Outcome of 7-day incubation with brain-derived Aβ-seeds isolated from patients with CAA on PLD3 levels in iPSC-neurons. Data presented are normalized to GADPH (mean ± S.E.M., *n=3*). \**p* < 0.05, \*\**p* < 0.01, \*\*\**p* < 0.001 by one-way ANOVA compared to nontreated cells. G) PLD3 ubiquitination is altered in AD as shown by representative confocal microscopy images of complexes between PLD3 and ubiquitin *(in situ PLA, red clusters)* in non-AD *(left panel)* and AD *(right panel)* sections, *n=7*. Data for quantification of PLA-puncta PLD3-ubiquitin complexes *(per field)* is presented as mean ± S.E.M., *n=7* with five images per case. \*\*\**p* < 0.001 by unpaired *t*-test. Nuclei were stained with TO-PRO^TM3^ *(blue)* and AD hallmarks with MX04 *(green).* The scale bars are 50 μm. H) The subcellular location of PLD3-ubiquitin complexes is non-lysosomal. The correlation coefficient of between PLA-puncta indicating PLD3-ubiquitin complexes and CTSD immunostaining is anti-correlated. Pearson correlation coefficient *(PCC, r)* is presented as mean ± S.E.M., *n=5*. The scale bars are 50 μm.

PLD3 was observed mainly in grey matter and was sparse in white matter (Fig. S3A). In non-AD grey matter, the PLD3 signal colocalized primarily with a neuronal marker (Fig. S3B-C) and occasionally with a microglia marker (Fig. S3D), but not with a marker for astrocytes (Fig. S3E). PLD3 was present in white matter, with more frequent co-localization with a microglia marker, but PLD3 level in AD white matter was not significantly different compared to non-AD white matter (Fig. S3F-G).

We evaluated the PLD3 levels by western blot in grey matter (Fig. 1C and Fig. S4A) and white matter (Fig. 1D and Fig. S4B) in homogenates of human temporal lobe tissue. We observed additional bands to the previously reported primary 50 and 70 kDa bands (42). These additional bands could be products of posttranslational modifications and were included in the total PLD3 quantification as they appeared to be specific in our verification experiments (Fig. S1). In grey matter, there was approximately a 38% reduction in total PLD3 levels in AD (Fig. 1C and Fig. S4A). No significant differences in total, lumenal, and membrane-associated PLD3 levels were found in white matter (Fig. 1D and Fig. S4B).

An initial report that PLD3 may regulate β-amyloid and/or APP levels was not reproducible in our work (21) and the work of others (23). Still, it remains unclear whether PLD3 levels are affected by β-amyloid. We evaluated the effect of a 7-day exposure of synthetic Aβ_1-42_ (1 µg/mL) on PLD3 levels in a human neuroblastoma cell line (SH-SY5Y cells, Fig. 1E and Fig. S5A). We detected a significant decrease in PLD3 on the third day but not in the following days (Fig. 1E). A substantial reduction in PLD3 level was found in analogous experiments using iPSC-derived neurons *(a cell line from a neurologically normal, young female donor)* incubated with brain-derived β-amyloid seeds from patients with cerebral amyloid angiopathy (CAA) (100 ng/mL) (Fig S5B-D). This decrease persisted after the third day (Fig. 1F and Fig. S5E). The average lysosomal size was increased after exposure to brain-derived β-amyloid seeds Fig. S6.

The decreased level of PLD3 in the brain is partly due to reduced transcription (43). Still, ubiquitination could contribute to accelerated degradation and/or altered localization within the brain, as was suggested in previous cell-culture based experiments (42). For this reason, we sought to visualize ubiquitinated PLD3 in human brain tissue. Ubiquitin immunochemistry revealed the marked accumulation of ubiquitin which is expected in AD brains (Fig. S7A-B). *In-situ* PLA confirmed proximity between PLD3 and ubiquitin, consistent with ubiquitin modification of PLD3 (Fig. 1G and Fig. S8C). In non-AD brain sections, PLA puncta corresponding to PLD3-ubiquitin complexes were sparse (Fig. 1G, *right panel*), while significantly greater levels of PLA puncta were observed in AD brain sections (Fig. 1G, *left panel*). PLD3-ubiquitin complexes were partially localized to tau pathology (Fig. 1G, *left panel*, and Fig. S8C). There was a 4.8-fold increase in the number of PLA puncta/field in AD compared to non-AD (Fig. 1G). We found scarce PLA puncta in technical and biological controls for *in-situ* PLA (Fig. S8D).

We combined *in-situ* PLA with IHC to visualize the subcellular location of PLD3-ubiquitin complexes. In both non-AD and AD brain sections, PLD3-ubiquitin complexes were negatively correlated with lysosome staining as shown by CTSD immunostaining (*p = 0.3149* by unpaired *t*-test with Welch’s correction, Fig. 1H), suggesting the lysosomal localization of PLD3 is lost when it is ubiquitinated.

### Loss of PLD3 alters lysosome morphology and function and remodels the lysosomal proteome

We obtained and validated PLD3 knockout cell lines, clone 3 (PLD3^−/−^) and clone 14 (PLD3^−/−*c14*^), in the SH-SY5Y neuroblastoma cell line, as shown in Fig. S1.

Lysosomal attributes in these cells, including size, density, and confluency, were characterized by transmission electron microscopy (TEM) (Fig. 2A). The size of individual lysosomes in PLD3^−/−^ cells was enlarged by 48% compared to WT cells (Fig. 2A). Lysosomal density and confluency, expressed by the number of lysosomes per unit area and the percentage of occupied area, respectively, were also increased in the PLD3^−/−^ lines (Fig. 2A).

**Figure 2.**
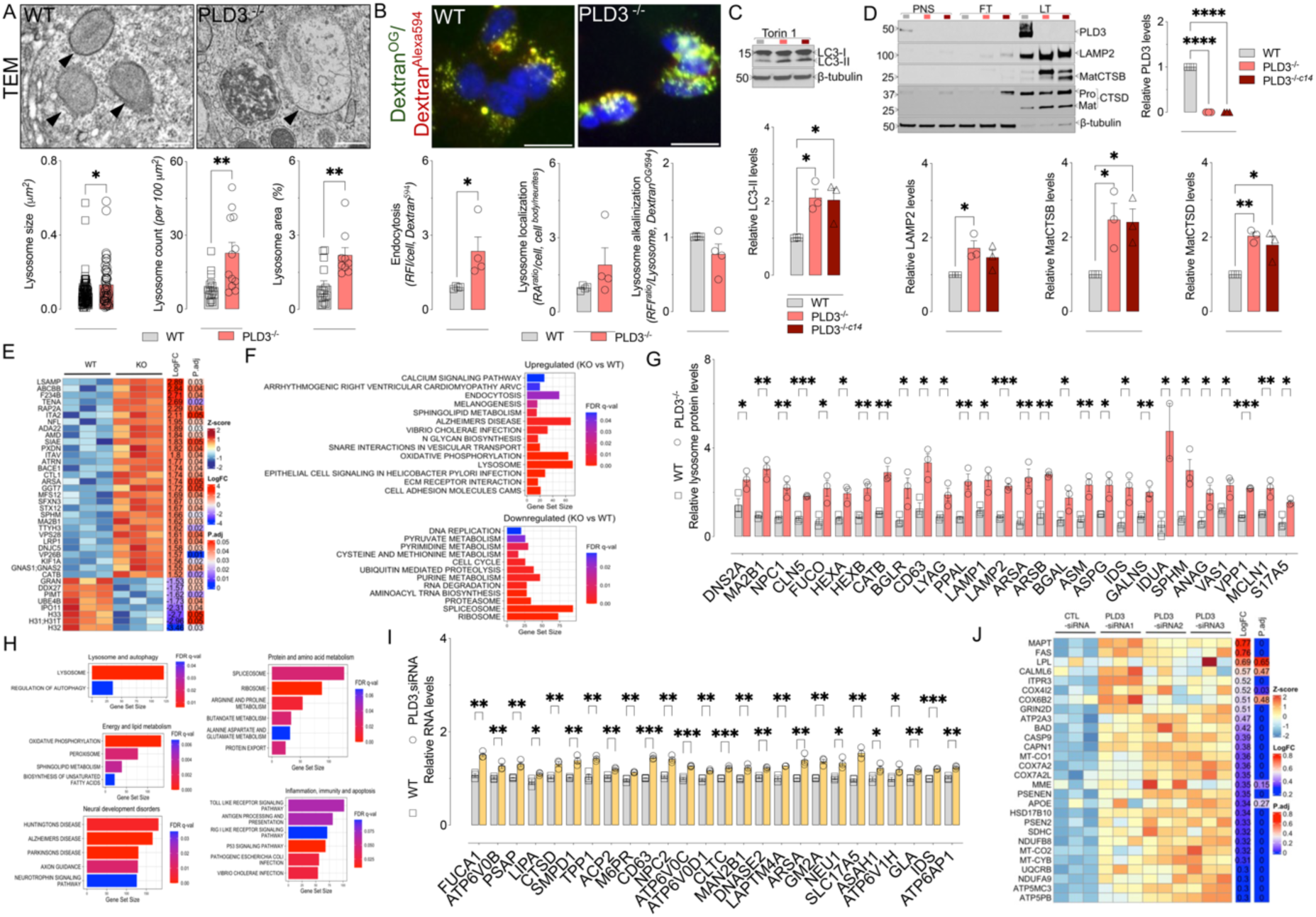
PLD3 deficiency alters the morphology and function of lysosomes. A) Quantification and comparison of lysosomal features *(size, density, and confluence)* between WT and PLD3*^−/−^* SHSY5Y cells by electron microscopy (TEM). Data presented mean ± S.E.M. For size: WT, *n=92* lysosomes; PLD3*^−/−^*, *n=51* lysosomes (\**p* < 0.05 by Welch’s unequal variances *t*-test). For lysosome density (count per 100 μm^2^): WT, *n=15* images; PLD3*^−/−^*, *n=12* images (\*\**p* < 0.01 Welch’s unequal variances *t*-test). For lysosome count (occupied area): WT, *n=15* lysosomes; PLD3*^−/−^*, *n=9* lysosomes (\*\**p* < 0.01, Welch’s unequal variances *t*-test). The scale bars are 500 nm. B) We evaluated lysosomal features of lysosomal function *(endocytic activity, alkalinization, and positioning)* between WT and PLD3*^−/−^* cells incubated with Dextran^OG^ *(pH-sensitive)* and dextran^Alexa594^ *(pH-insensitive)*. Data presented mean ± S.E.M., *n=4*. \**p* < 0.05 by unpaired *t*-test. The scale bars are 50 µm. C) Western blot analysis of LC3-I and LC3-II in WT and PLD3*^−/−^* cells treated with Torin 1 (1 µM) for 3 hours. Proteins were quantified, expressed relative to β-tubulin, and compared between phenotypes. Data presented means ± S.E.M., *n=3*. \**p* < 0.05 by one-way ANOVA followed by Dunnett’s multiple comparisons test. D) Lysosomal protein levels in lysosomes isolated from WT and PLD3*^−/−^*cells. Data presented normalized to β-tubulin mean ± S.E.M., *n=3*. \**p* < 0.05, \*\**p* < 0.01, and \*\*\*\**p* < 0.0001 by one-way ANOVA followed by Dunnett’s multiple comparison test. PNS: post-nuclear supernatant. FT: flow-through. LF: lysosome fraction. E) Proteomic analysis of the lysosomal lysates from WT and PLD3*^−/−^* cells. The differentially expressed proteins were represented by a heatmap (*n=3*, minimum change in magnitude ≥ 1.5-fold change, false discovery rate (FDR) less than 5% (< 0.05). F) Normalized proteomic data were subject to gene set enrichment analysis (GSEA), and the differentially regulated pathways in PLD3*^−/−^* versus WT were summarized and represented by bar plots (FDR q value < 0.05). G) Selected lysosomal proteins from the proteomic analysis were quantified and compared between WT and PLD3*^−/−^*, and the differentially regulated proteins were represented by bar graphs (mean ± S.E.M., *n=3*, multiple unpaired *t*-tests). H) RNA-Seq data underwent GSEA. Differentially enriched pathways in PLD3_siRNAs *(n=9)* versus CTL_siRNA were summarized and represented by bar plots (FDR q value < 0.05). I) RNA expression for genes related to lysosome pathways was calculated and compared between CTL_siRNA *(n=3)* and PLD3_siRNAs *(n=9)*, and significantly different genes were represented by bar graphs (mean ± SEM, multiple unpaired t test). J) RNA expression in genes related to AD was calculated and compared between CTL_siRNA *(n=3)* and PLD3_siRNAs *(n=9)*, and significantly different genes were represented by a heatmap (mean ± SEM, multiple unpaired t test).

Relative lysosomal pH was evaluated using the ratio of two fluorophores, Oregon Green (pH-sensitive) and Alexa^594^ (pH-insensitive), which were trafficked to lysosomes by conjugation to dextran (Fig. 2B and Fig. S8). Dextran localization to lysosomes was confirmed by colocalization and correlation analysis between the dextran signal and LAMP1-RFP (Fig. 2B and Fig. S8A). We did not observe a significant change in the pH of lysosomes in PLD3^−/−^ cells (Fig. 2B and Fig. S8A). Lysosome size increased by 61% (in line with the TEM result), and confluency also increased in this fluorescence assay (Fig. S8A). Based on lysosomal labelling with dextran, lysosome density was unchanged (Fig. S8A). Lysosome-related cellular processes, such as endocytosis and autophagy, were enhanced by 2.5-fold and 2-fold respectively in PLD3^−/−^cells (Fig. 2B-C).

To assess changes in the lysosomal proteome, we isolated lysosomes from WT and PLD3^−/−^ cells using dextran-coated superparamagnetic iron oxide (SPIO) nanoparticles and magnetic chromatography, followed by label-free proteomic studies on the lysates of lysosomal fractions. Cathepsin D (CTSD), cathepsin B (CTSB), and LAMP2 were highly enriched in the lysosomes of PLD3^−/−^ cells as compared to WT cells (Fig. 2D), while cytosolic-proteins like β-tubulin were appropriately absent in the lysosomal fraction (Fig. 2D).

Proteomic analysis of lysosomal lysates from each cell line identified 3,075 proteins, of which 81 were significantly upregulated and 72 were downregulated in PLD3^−/−^ lines compared to WT cells (Fig. 2E). Gene set enrichment analysis (GSEA) revealed that signaling pathways related to lysosomal biogenesis, endocytosis, and calcium signaling were significantly enriched in PLD3^−/−^compared to WT cells (Fig. 2E-G and Fig. S9). PLD3^−/−^ cells were also enriched in vesicle trafficking proteins, such as SNARE (Soluble NSF Attachment Protein Receptor) proteins. We also observed that proteins related to sphingolipid metabolism and proteins relevant to AD were differentially enriched between WT and PLD3^−/−^ cells (Fig. 2G).

We then examined the impact of PLD3 deficiency on the cellular transcriptome. Total RNA from WT and PLD3^−/−^ cells underwent RNA sequencing, followed by differential gene expression (DGE) analysis and gene set enrichment analysis (GSEA). The sequencing data was summarized into 60,662 genes, of which 7,673 genes were differentially expressed between WT and PLD3^−/−^cells, with 4,015 upregulated and 3,658 downregulated (FDR<0.05; (Fig. S10). The upregulated genes were primarily associated with signaling pathways involved in cell-cell interaction, neural development, and inflammatory conditions (Fig. S10). Signaling pathways related to lysosomal biogenesis, endocytosis, and AD were also upregulated in PLD3^−/−^ cells compared to WT cells. However, their significance did not reach the FDR<10% threshold (Fig. S10).

To explore the immediate transcriptomic changes associated with the milder phenotype resulting from PLD3 deficiency, PLD3 was knocked down in SH-SY5Y cells using siRNA, followed by RNA sequencing and analysis, and verification through western blot (Fig. S11). Compared to control siRNAs (CTL), all three PLD3 siRNAs achieved 60-90% knockdown of PLD3 (Fig. S11). DGE analysis revealed 2,830 genes that were differentially expressed between PLD3 KD and CTL cells (Fig. S11). 2,118 genes were upregulated, and 712 genes were downregulated in PLD3 KD cells compared to CTL cells (FDR<0.05; Fig. S11). We also observed significant enrichment in signaling pathways associated with lysosomes and autophagic flux, such as lysosome biogenesis, endocytosis, mTOR signaling, and autophagy signaling, in PLD3 KD cells relative to CTL cells (Fig. 2H). Additionally, signaling pathways previously reported to be associated with PLD3 or lysosomes were also differentially enriched between PLD3 KD cells and CTL cells (Fig. 2H). The transcripts associated with lysosome biogenesis and AD pathologies are illustrated in Fig. 2I and 2J, showing a significant increase in PLD3 KD cells compared to CTL cells.

### Loss of PLD3 increases CTSB level and activity and alters its subcellular location

Because both synthetic and human brain-derived β-amyloid seeds decrease PLD3 levels, and PLD3^−/−^ in SH-SY5Y cells altered the level of some lysosome resident protein, particularly CTSB, we evaluated CTSB levels in WT and PLD3^−/−^ cells exposed to synthetic Aβ_1-42_ (1 μg/ml) for 1, 3, 5, and 7 days (Fig. 3A). Mature CTSB (MatCTSB) levels were increased in WT and PLD3^−/−^ cells after exposure to Aβ_1-42_. This increase was more substantial and sustained in PLD3^−/−^ cells (Fig. 3A). Additionally, we found that PLD3^−/−^ cells had increased specific CTSB activity compared to WT cells (Fig. 3B). However, increased CTSB activity did not lead to improved lysosomal degradative capacity as measured by DQ assay (Fig. 3C). PLD3^−/−^ cells treated with β-amyloid particles isolated from human brain had an accumulation of intracellular β-amyloid compared to wild type cells after a three-day incubation. Similarly, human synthetic alpha-synuclein accumulated in PLD3^−/−^ cells compared to wild type cells (Fig. 3C). PLD3^−/−^ cells were able to degrade synthetic Aβ_1-42_ to the same extent as WT (Fig. 3C).

**Figure 3.**
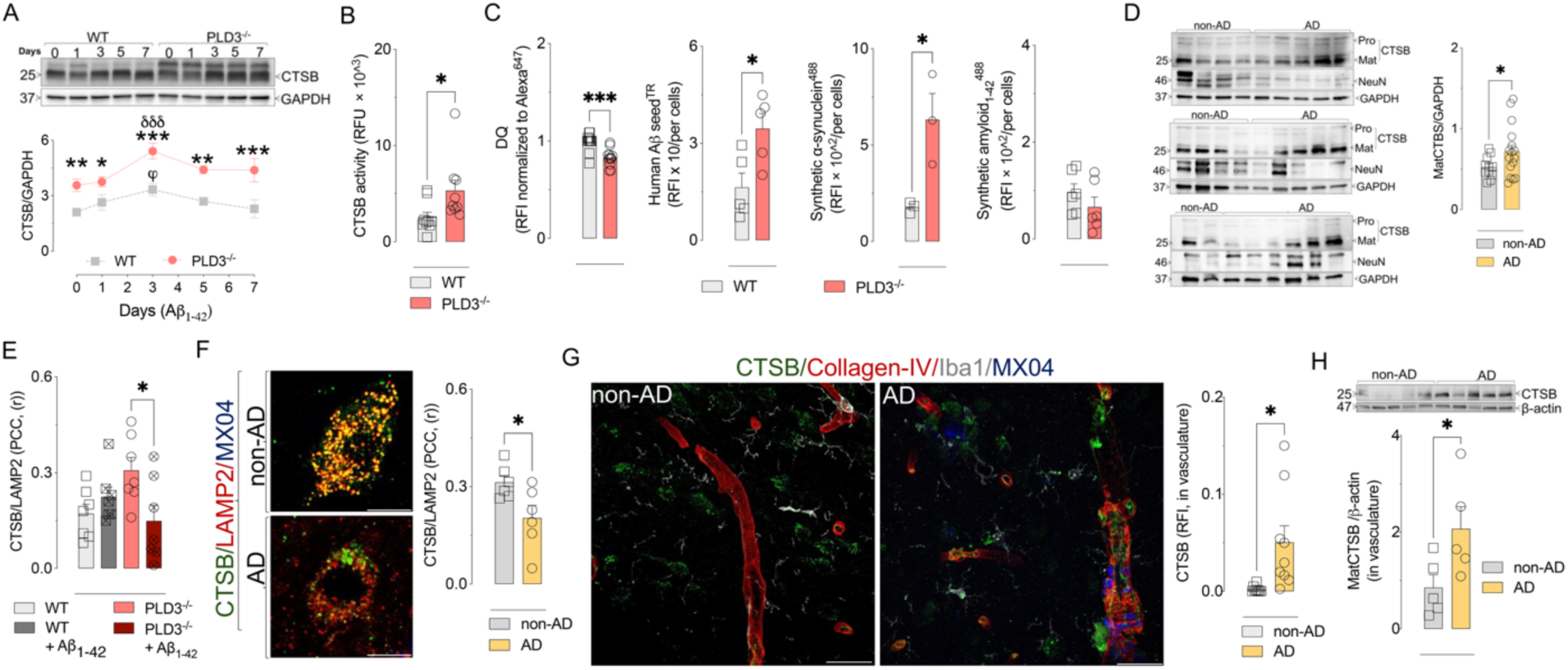
Loss of PLD3 triggers changes in CTSB level, activity, and subcellular location. A) Outcome of long-term incubation with synthetic Aβ_1-42_ (1 μg/mL) on CTSB levels in WT and PLD3^−/−^ cells. Data presented mean ± S.E.M., *n=4*. \**p* < 0.05, \*\**p* < 0.01, and \*\*\**p* < 0.001 by two-way ANOVA compared to WT.^ϕ^*p* < 0.05 by one-way ANOVA compared to basal WT cells. ^δδδ^*p* < 0.001 by one-way ANOVA compared to basal PLD3^−/−^ cells. B) Enzymatic CTSB activity in PLD3^−/−^ cells compared to WT. Data normalized to enzymatic activity in the presence of the potent and selective inhibitor of CTSB, CA074 (1 μM). Data presented mean ± S.E.M., *n=9*. \**p* < 0.05 by unpaired *t*-test. C) Intracellular accumulation of protein aggregates in WT and PLD3^−/−^ cells. Data presented mean ± S.E.M., *n=12 (for DQ), n=5 (for human Aβ seed^TR^), n=3 (for human synthetic α-synuclein^488^), and n=6 (for human synthetic Aβ*_1-42_*^488^).* \**p* < 0.05 and \*\*\**p* < 0.001 by unpaired *t*-test. D) Western blot analysis of CTSB in the superior temporal lobe protein fraction. Each lane corresponds to a donor of the indicated group. Data presented mean ± S.E.M., non-AD *(n=10),* and AD brains *(n=16)*. *\*p* < 0.05 by Welch’s unequal variances *t*-test. E) Immuno-colocalization of CTSB and LAMP2 in WT and PLD3^−/−^ cells with or without synthetic Aβ_1-42_ incubation. Bars are mean ± S.E.M., *n=7*. \**p* < 0.05 by one-way ANOVA followed by Tukey’s multiple comparisons test. F) Representative confocal microscopy images of CTSB-LAMP2 colocalization in non-AD and AD sections. Data presented are means (Pearson’s correlation coefficient, PCC, (r)) ± S.E.M., *n=7*, each with five images \**p* < 0.05 by unpaired *t*-test. The scale bars are 5 μm. G) CTSB in the cerebral vasculature. Representative confocal microscopy images and quantification of CTSB expression pattern *(green)* in the temporal lobe from non-AD and AD cases. Mean CTSB intensity in vascular regions was quantified solely in the collagen-IV immunoreactive area *(red)*. β-amyloid aggregates and cerebrovascular amyloid deposition were detected with MX04 *(blue)* and microglia by the marker Iba1*(grey)*. Data presented mean ± S.E.M., *n=9*, each with at least five images. \**p* < 0.05 by unpaired *t*-test. The scale bars are 50 μm. H) Western blot analysis of the level of CTSB in isolated human vessels from the temporal lobe protein fraction from non-AD and AD cases. Bars are mean ± S.E.M., *n=5*. \**p* < 0.05 by unpaired *t*-test.

In human brain homogenates of superior temporal gyrus grey matter, pro-cathepsin B (ProCTSB) and mature cathepsin B (MatCTSB) were detectable at ∼43-46 kDa and 25-26 kDa, respectively (Fig. 3D and S14). We found an increase in MatCTSB level in AD compared to non-AD cases when levels were normalized to the neuronal marker NeuN or to GADPH (Fig. 3D and S13).

Since a change in subcellular CTSB location may also contribute to the deficit in the hydrolysis of protein aggregates despite increasing overall CTSB level and activity, we measured the colocalization of CTSB with lysosomal marker LAMP2 in the cell model and in human brain tissue. We found a reduction of CTSB co-localization with LAMP2 in PLD3^−/−^ cells incubated with Aβ_1-42_ (1 μg/ml) for 7-days compared to untreated PLD3^−/−^ cells, but not in treated WT cells compared to untreated WT cells (Fig. 3E). Likewise, a significant portion of CTSB is not colocalized to lysosomes in AD brain tissue but is instead localized in the extra-lysosomal compartment (Fig. 3F and Fig. S14A-B). This extracellular localization included the human brain microvasculature. We stained CTSB and collagen-IV; CTSB staining was present in the vasculature in control cases but was much more abundant in AD cases, especially in those with CAA (Fig. 3G and Fig. S14C-D). To confirm this association, we isolated human vessels from the superior temporal lobe and measured CTSB levels, revealing an increase in CTSB in AD compared to non-AD (Fig. 3H).

### PLD3 regulates TFEB/TFE3 degradation through the proteasome rather than lysosomes

Based on our proteomic analysis of isolated lysosomes from PLD3^−/−^ cells, we hypothesized that PLD3 might play a role in lysosomal biogenesis. We assessed mRNA levels of TFEB and TFE3 and found no significant difference between WT and PLD3^−/−^ cells, suggesting that any potential regulation of TFEB and TFE3 by PLD3 does not occur at the transcriptional level (Fig. S15A). In contrast, TFEB and TFE3 protein levels were markedly elevated in PLD3^−/−^ cells compared to WT cells under a range of conditions including no stimulation (NS), dextran treatments to induce endocytosis, nutrient starvation (HBSS), and Torin 1 treatment to induce autophagy (Fig. 4A and Fig. S15B). Exogenous TFEB, expressed as GFP N-terminal tagged TFEB, also showed increased expression in PLD3^−/−^ cells versus WT cells (Fig. 4B). We found that in both PLD3 knockdown (KD) and in PLD3^−/−^ cells, the expression trends in TFEB and TFE3 were similar (Fig. S15C). Consistent with these results, the overexpression of PLD3 in HeLa cells led to a significant reduction in TFEB and TFE3 levels (Fig. 4C). Interestingly, AD-associated variants in PLD3, e.g., V232M or K486R, did not alter its effects on TFEB metabolism (Fig. 4C). These data suggest that PLD3 is involved in TFEB/TFE3 protein metabolism, and PLD3^−/−^ leads to their accumulation.

**Figure 4.**
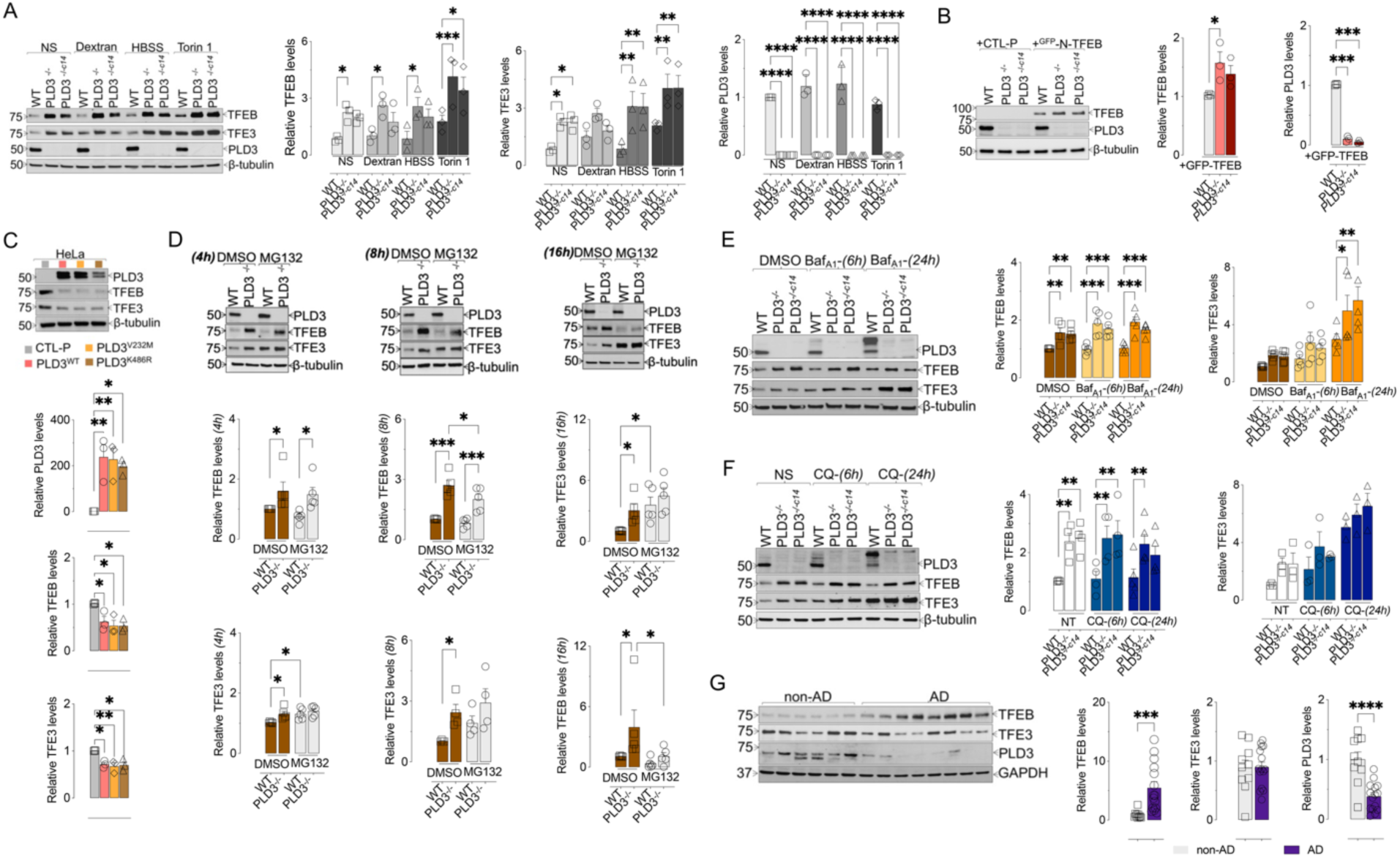
PLD3 regulates TFEB/TFE3 metabolism. The proteasome, rather than lysosomes, is predominantly involved in PLD3-mediated TFEB/TFE3 metabolism. A) Western blot analysis of TFEB, TFE3, and PLD3 in WT and PLD3*^−/−^* cells treated with vehicle (NS, *non-stimulation*), dextran (0.25 mg/mL), HBSS, or Torin 1 (1 mM) for 3 hours. Proteins were quantified, expressed compared to β-tubulin, compared between genotypes, and presented as mean ± S.E.M., *n=3*. \**p* < 0.05, \*\**p* < 0.01, and \*\*\*\**p* < 0.0001 by two-way ANOVA followed by uncorrected Fisher’s LSD. B) Western blot analysis of TFEB and PLD3 in WT and PLD3*^−/−^* cells transfected with control plasmids (CTL-P) or GFP-tagged TFEB plasmids (GFP-N-TFEB) for 2 days. TFEB and PLD3 were quantified relative to β-tubulin and compared between genotypes (mean ± S.E.M., *n=3*). \**p* < 0.05, and \*\*\*\**p* < 0.0001 by two-way ANOVA followed by uncorrected Fisher’s LSD. C) Western blot analysis of TFEB, TFE3, and PLD3 in HeLa cells transfected with control plasmids (CTL-P) or PLD3 plasmids for 2 days. Proteins were quantified relative to β-tubulin and compared between genotypes (mean ± S.E.M., *n=3*). \**p* < 0.05, and \*\**p* < 0.01 by one-way ANOVA followed by Dunnett’s multiple comparison test. D) Western blot analysis of TFEB, TFE3, and PLD3 in WT and PLD3*^−/−^* cells treated with MG132 (1 mM) for 4, 8, and 16 hours. Proteins were quantified relative to β-tubulin and compared among experimental conditions (mean ± S.E.M., *n=4-5*). \**p* < 0.05, \*\*\**p* < 0.001 by two-way ANOVA followed by uncorrected Fisher’s LSD. E and F) Western blot analysis of TFEB, TFE3, and PLD3 in WT and PLD3*^−/−^* cells treated with (E) bafilomycin A1 (BafA1, 200 nM, *n=5*) or (F) chloroquine (CQ, 50 mM, *n=4*) for 6 hours and 24 hours. Proteins were quantified relative to β-tubulin and compared among experimental conditions (mean ± S.E.M). \**p* < 0.05, \*\**p* < 0.01, and \*\*\**p* < 0.001 by two-way ANOVA followed by uncorrected Fisher’s LSD. G) Representative western blot analysis of TFEB, TFE3, and PLD3 levels in superior temporal lobe protein lysate from human brain. Each lane corresponds to a donor of the indicated group. Data are normalized to GADPH (means ± S.E.M., non-AD *n=10*, and AD *n=15*). \*\*\**p* < 0.001 by Welch’s unequal variances *t*-test.

We investigated the potential involvement of the proteasome and/or lysosome in PLD3 regulation of TFEB/TFE3 degradation. Treatment with MG132, a proteasome inhibitor, caused a significant accumulation of TFE3 and resulted in comparable levels of TFE3 between WT and PLD3^−/−^ cells at all three time points (Fig. 4D), suggesting that PLD3 facilitates TFE3 degradation via the proteasome. In contrast, TFEB levels declined over time following MG132 treatment, indicating that the proteasome was indirectly involved in PLD3-mediated TFEB metabolism (Fig. 4D). Treatment with another proteasome inhibitor, bortezomib, produced similar effects on TFEB/TFE3 (Fig. S16A). To focus on protein metabolism, we blocked protein translation using cycloheximide (CHX) and assessed TFEB/TFE3 levels following MG132 treatment. MG132 generated similar effects on TFEB/TFE3 with the arrest of protein translation, supporting the conclusion that the proteasome is involved in PLD3-mediated TFEB/TFE3 protein metabolism (Fig. S16A). Interestingly, CHX treatment led to a sharp decline in TFE3 levels but a gradual decrease in PLD3 and TFEB levels, suggesting that TFE3 is a short-lived protein, while TFEB and PLD3 are long-lived proteins (Fig. S17A). Immunoprecipitated (IP) TFE3 showed heightened ubiquitination in WT cells versus PLD3^−/−^ cells upon MG132 treatment (Fig. S17B). Concerning the potential involvement of lysosomes in TFEB/TFE3 metabolism, treatments with lysosome alkalinizing agents, chloroquine (CQ) or bafilomycin A1 (BAF), exhibited no significant effects on TFEB/TFE3 levels, which suggests that these proteins are not predominantly degraded via lysosomal pathways (Fig. 4E and 4F). Nevertheless, we did observe an accumulation of TFE3 after prolonged exposure to CQ or BAF. This could be due to the induction of lysosomal stress following CQ or BAF exposure, which activated TFE3 signaling rather than directly impacting TFE3 protein metabolism (Fig. 4E and 4F).

TFEB levels in the brains of AD patients were significantly higher than in non-AD patients (Fig. 4G). However, this pattern was not observed for TFE3 (Fig. 4G).

### Loss of PLD3 enhances nuclear translocation and transcriptional activity of TFEB/TFE3

To determine whether the change in TFEB/TFE3 protein levels in PLD3^−/−^ cells altered TFEB/TFE3 signaling, we assessed TFEB/TFE3 levels in the cytoplasmic *(c)* and nuclear *(n)* compartments using western blotting. Under basal conditions and conditions known to induce autophagy, including nutrient deprivation (HBSS) or mTOR inhibition (Torin 1), the protein levels of TFEB/TFE3 in the nuclear compartment of PLD3^−/−^ cells were much higher than those in WT cells (Fig. 5B). However, we did not find a significant difference in the protein ratio of nuclear TFEB/TFE3 to cytosolic TFEB/TFE3 between WT and PLD3^−/−^ cells (Fig. 5B). Surprisingly, we observed some nuclear localization of PLD3 in WT cells (Fig. 5A).

**Figure 5.**
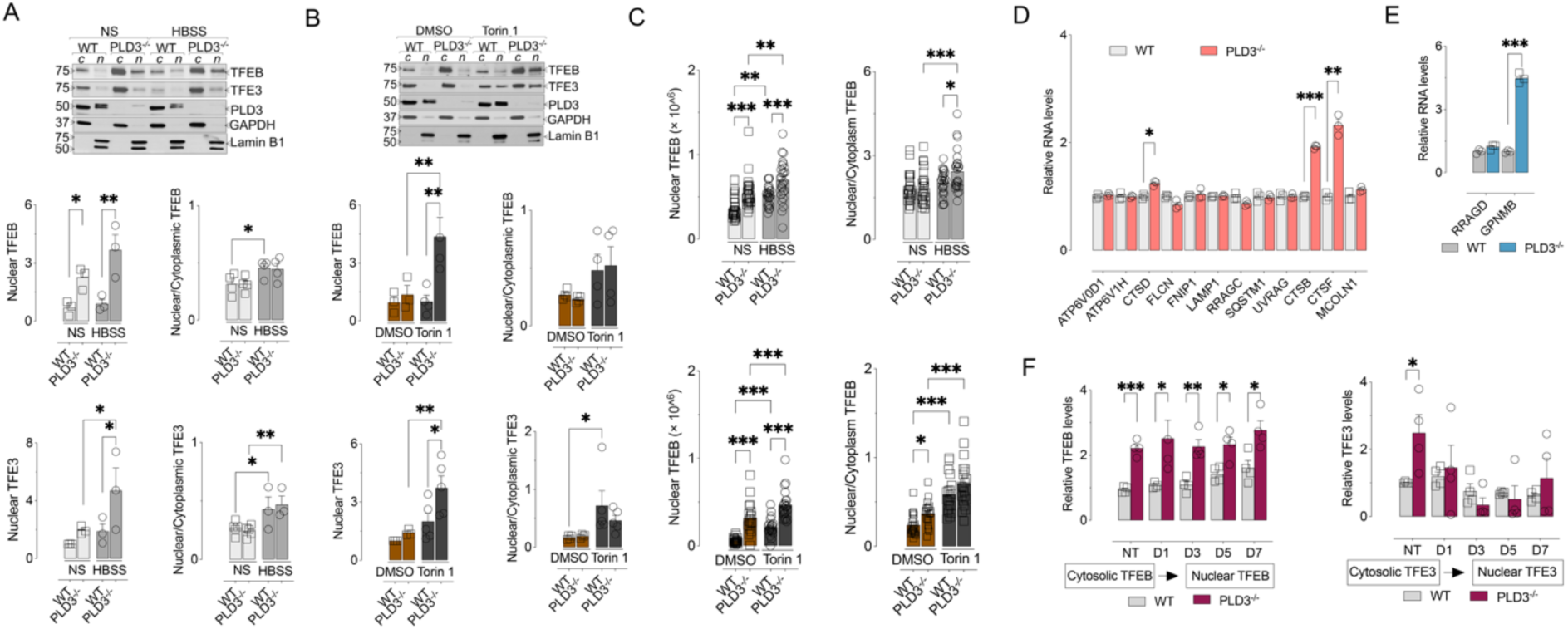
Loss of PLD3 augments TFEB/TFE3 transcription and nuclear localization. A and B) Western blot analysis of TFEB, TFE3, and PLD3 in nuclear extract *(n)* and cytoplasmic extract *(c)* of WT and PLD3*^−/−^* cells treated with (A) HBSS (*vs.* NS *(non-stimulation)*), or (B) 1 μM Torin 1 (*vs.* DMSO) for 3 hours. Proteins were quantified and normalized to GAPDH *(cytoplasmic)* or Lamin B1 *(nuclear)*. Proteins in *n* or their *n*-to-*c* ratios were compared among experimental conditions and presented as mean ± S.E.M., *n=3-5*. \**p* < 0.05 and \*\**p* < 0.01 by two-way ANOVA followed by uncorrected Fisher’s LSD. C) Immunostaining analysis of TFEB location in WT and PLD3*^−/−^* cells treated with HBSS (*vs.* NS *(non-stimulation)*) or 1 μM Torin 1 (*vs.* DMSO) for 3 hours. Data presented means ± S.E.M., *n=20-25* fields. \**p* < 0.05, \*\**p* < 0.01, and *** *p* < 0.001 by two-way ANOVA followed by uncorrected Fisher’s LSD. D and E) Transcript levels of genes containing the CLEAR motif (D) or genes responsive to TFEB (E) in WT and PLD3*^−/−^* cells were calculated and compared between WT and PLD3*^−/−^*. Data presented means ± S.E.M., *n=3*. \**p* < 0.05, \*\**p* < 0.01, and *** *p* < 0.001 by multiple unpaired *t-*tests. F) Outcome of long-term incubation with synthetic Aβ_1-42_ on TFEB and TFE3 location. Proteins were quantified relative to β-tubulin and compared among experimental conditions (mean ± S.E.M., *n=4*). \**p* < 0.05, \*\**p* < 0.01, and *** *p* < 0.001 by two-way ANOVA followed by Fisher’s LSD multiple comparison test.

Immunocytochemical staining of TFEB confirmed these findings, showing increased TFEB levels and nuclear translocation in PLD3^−/−^ cells compared to WT cells (Fig. S18). Exogenous TFEB, expressed as GFP N-terminal tagged TFEB, also exhibited increased nuclear translocation in PLD3^−/−^ cells versus WT cells, with or without Torin 1 treatment (Fig. S18). Corresponding to the increased nuclear translocation of TFEB/TFE3, the transcription of their targeted genes, which contain coordinated lysosomal expression and regulation (CLEAR) motif, was also augmented (Fig. 5D) in PLD3^−/−^ cells compared to WT cells. Moreover, the mRNA levels of two genes, RRAGD and GPNMB, known to respond to TFEB, were upregulated (Fig. 5E).

Next, we examined TFEB/TFE3 signaling in neuroblastoma cells after exposure to synthetic Aβ_1-42_ at various intervals. In WT and PLD3^−/−^ cells, Aβ_1-42_ exposure mediated TFEB and TFE3 nuclear translocation from day 3 through day 7 (Fig. 5F). The nuclear translocation of TFEB and TFE3 in response to Aβ_1-42_ exposure was much higher in PLD3^−/−^ cells than in WT cells (Fig. 5F).

## Discussion

This study links PLD3 to lysosome abnormalities in AD. PLD3 levels were reduced in human AD by 38% compared to non-AD human brains. In cultured cells, PLD3 deficiency resulted in changes in lysosomal morphology, biogenesis, and degradative function, while lysosomal acidification remained relatively intact. These findings are significant given the pivotal role of the endosomal and lysosomal systems in proteinopathies.

We and others previously described the distribution of PLD3 in histological staining of human brain tissue, reporting that it is primarily in neuronal lysosomes and enriched in dystrophic neurites around neuritic plaques (20, 21). In the current study, we further characterized the level and distribution of PLD3 in white versus grey matter across multiple brain regions to refine our understanding of its association with the neuropathologies of AD. Our observations in cell models suggest that Aβ exposure may contribute to the reduced level of PLD3 we observed in neurons. In line with diminished PLD3 in AD cortex, ubiquitination of PLD3 was markedly increased in AD tissue. PLD3-ubiquitin complexes were primarily non-lysosomal and frequently colocalized with neurofibrillary tangles and neuropil threads. This is consistent with a previous finding from a cellular model system showing that ubiquitinated PLD3 fails to traffic to the lysosome (42).

Cultured neuroblastoma (SH-SY5Y) cells with PLD3 deletion had numerous lysosomal abnormalities including lysosomal enlargement and decreased efficiency in processing aggregated α-synuclein and β-amyloid particle derived from human brain tissue. There was no difference in the cells’ ability to degrade synthetic Aβ_1-42_, but this may be due to differences in aggregate size, endocytosis rate, or a number of other factors. There was a general pattern of impaired degradative function in PLD3^−/−^ lysosomes. In line with this result, we found the proteolytic capacity of the cells was compromised based on the DQ assay. In contrast with these findings, we found that the enzymatic activity of CTSB and the level of both CTSD and CTSB were increased in PLD3−/− cells.

We isolated lysosomes from these cells using magnetic chromatography and performed ICP-MS to characterize the lysosomal proteome. We confirmed that the lysosomes also had elevated levels of CTSB, LAMP1 and 2, Hex A and B, β-galactosidase, and CD63, among numerous other lysosomal proteins. We also found that signaling pathways relevant to AD were enriched in PLD3^−/−^ lysosomes, including elevations in presenilin 2 and LRP1. This broad alteration in the proteome led us to hypothesize that PLD3 may alter proteolytic lysosome machinery and have a role in regulating lysosomal biogenesis.

We also discovered that in differentiated PLD3^−/−^ neurons stressed by prolonged exposure to β-amyloid, the fraction of CTSB associated with lysosomes was decreased, despite increased total levels of CTSB. We also observed extra-lysosomal CTSB in neurons and in extracellular matrices in human AD cortex. The loss of lysosomal localization of CTSB may account for the increase in CTSB activity in PLD3^−/−^ neurons, while total lysosomal degradative capacity was decreased. CTSB may be leaking from damaged/enlarged lysosomes into extra-lysosomal compartments. CTSB is primarily localized to neuronal lysosomes in non-disease conditions, but CTSB leakage from injured lysosomes can lead to apoptosis (4) and pyroptosis mediated by caspase-1(44). Recent studies have found that CTSB may be enzymatically active in extralysosomal locations, including the nucleus (45) and extracellular space (46–48), because, in contrast with other lysosomal proteases, CTSB retains proteolytic activity at neutral pH (49, 50). This activity at neutral pH is key to CTSB-mediated adverse effects in the cytoplasm (44, 51–54) and the degradation of matrix components in the extracellular space (55–57). Our results suggest that loss of PLD3 not only regulates CTSB abundance but may contribute to loss of its lysosomal localization.

We next confirmed that PLD3 has a role in regulating lysosomal biogenesis by demonstrating that TFEB was markedly elevated in the PLD3^−/−^ cells. TFEB and TFE3 are master regulators of lysosomal biogenesis and autophagy; their role as transcription factors for lysosome-related genes activates lysosome production (58–60). Unlike conventional lysosomal stressors, such as amino acid deprivation, the deletion of PLD3 did not significantly impact mTOR and calcineurin activity. The phosphorylation level of endogenous TFEB at Ser^142^ and Ser^211^ was undetectable in SH-SY5Y cells. However, we observed a marked increase in TFEB and TFE3 protein levels (but not mRNA levels), and a corresponding increase in nuclear TFEB/TFE3 levels upon PLD3 knockout, suggesting that PLD3 regulates TFEB/TFE3 metabolism at the protein level. We assessed whether PLD3 mediated the degradation of TFEB and TFE3 via the proteasome. While treatment with proteasome inhibitors increased TFE3 levels, TFEB levels significantly fell with proteasome inhibition.

Additionally, while ubiquitin-mediated degradation of TFEB via the proteasome has been reported to be initiated by phosphorylation of TFEB by the IKK complex (61), our study did not show significant changes in IKKα/β, and the level of phosphorylated TFEB was undetectable in our cellular models (Fig. S16C and D). We speculate that TFEB is regulated by one or more proteins metabolized through the proteasome, but further studies are needed to delineate the mechanisms by which the proteasome influences TFEB metabolism. Moreover, because PLD3 is primarily localized in lysosomes (9), it will be essential to identify the mediators that transmit signals from lysosomes to TFEB and TFE3.

## Conclusion

PLD3 has a regulatory role in neuronal lysosomal function and biogenesis. Loss of PLD3 reduces lysosomal degradative function and may permit cathepsin B trafficking to extra-lysosomal compartments. Loss of PLD3 also promotes neuronal biogenesis by increasing the key transcription factors TFEB and TFE3. Consequently, restoring neuronal PLD3 levels may be an experimental approach to improving neuronal proteostasis in AD.

## Acknowledgements

We thank the many patients and their families who have donated brain tissue to support this and other necessary studies. We also acknowledge the contributions of the Vanderbilt Cell Imaging Shared Resource (CISR) Core for transmission electron microscopy, the Vanderbilt Technologies for Advanced Genomics (VANTAGE) Core for RNA sequencing, and the Mass Spectrometry Research Center for proteomic analysis.

We also acknowledge the contributions of Drs. Ketaki A. Katdare and Ethan S. Lippmann for providing the IPSC-neurons cells.

## Author contributions

The authors confirm contribution to the paper as follows: study conception and design: W.R-F., Y.W., C.C-T and M.S.S. PLD3 expression, distribution, and post-translational modification in PLD3⁻/⁻cells and brain tissues; CTSB level, activity, and localization changes in PLD3⁻/⁻ cells and brain tissues; and assays for intracellular accumulation of amyloid and α-synuclein. W.R-F, C.C-T, S.J., E.W, A.P, T.A., A.K., A.S, N.H,. S.J.F, and V.A.J performed the corresponding experimental work and data collection, as presented in Figures 1 and 3 and Supplementary Figures 1, 2, 3, 4, 5, 7, 13, and 14. Analyses of lysosomal morphology, and function in PLD3⁻/⁻ cells or brain tissues; proteomic profiling of lysosomal proteins; whole-cell RNA-seq transcriptomic profiling and related computational analyses; and mechanistic studies elucidating the link between PLD3, amyloid and TFEB/TFE3 signaling and metabolism: Y.W., P.R., and N.A. performed the corresponding experimental work and data collection, as presented in Figures 2, 4, and 5 and Supplementary Figures 6, 8, 9, 10, 11, 12, and 15-18. W.R-F and MSS supervised the study. Analysis and interpretation of results, draft manuscript preparation, and substantial revision: W.R-F., Y.W., S.J., E.W., A.P., N.H., T.J.H., B.C.M., J.A.M., and M.S.S.

All authors contributed to the final revision of the manuscript and approved the version submitted for publication.

## Funding

This project is financially supported by grants from the National Institutes of Health (1R56AG074279, 1K76AG060001) and a National Institutes of Health training grant (5T32AG058524) to M.S.S, and a R35GM144112 to J.A.M.

## Data availability statement

The data supporting this study’s findings are available from the corresponding author upon reasonable request.

## Competing interests

MSS has received research support from the NIH (1RF1NS129735, R01AG078803, RF1NS130334, and R21AG070859), American Federation of Aging Research (K76AG060001/Beeson Scholar), the NeuroScience Innovation Foundation. He has received payments for consulting from Greycourt and Co, Inc, Guidepoint, Raymond James Financial, Inc, and has received payments for expert testimony in legal cases. He chairs the DSMB for the REVISION trial, serves on the medical advisory board for Septagen and is the research integrity officer for the Journal of Alzheimer’s Disease. TH serves on the scientific advisory board for Circular Genomics and serves as the Deputy Editor for Alzheimer’s & Dementia: TRCI The rest of authors declare no competing interests.

## Notes

### Summary of Updates

This updated version includes a demographics table to describe the human tissue samples. Additionally, during pre-submission data quality checks, we identified one human brain sample that was grouped incorrectly – this has been corrected in the affected figures and the resulting quantifications have been updated. The key conclusions are unaffected.

