## Supplementary material for "Phospholipase D3 regulates lysosomal morphology, biogenesis, and function in neurons"

**Table 1.** Samples demographics de-identified neuropathological information.

| Age at death / Sex | Primary clinical categorization | AD neuropathology | CAA grade | PMI | Other neuropathologies |
|---|---|---|---|---|---|
| 71/M | Alzheimer's disease | Severe (A3, B3, C3) | Severe CAA | 12h | Right parietal hemorrhage |
| 81/M | Alzheimer's disease | Low (A2, B1, C2) | Severe CAA | 19h | Hemorrhages |
| 81/F | Alzheimer's disease | Severe (A3, B3, C3) | Severe CAA | 35h53' | None |
| 68/F | Alzheimer's disease | Severe (A3, B3, C3) | Severe CAA | 9h | None |
| 68/F | Alzheimer's disease | Severe (A3, B3, C3) | Low CAA | 15h | None |
| 70/M | Alzheimer's disease | Severe (A3, B3, C3) | Severe CAA | 11h | Tau in middle and superior temporal gyrus |
| 61/M | Alzheimer's disease | Moderate-severe (A2, B2, C3) | Moderate-severe CAA | 10h | None |
| 73/M | Alzheimer's disease | Moderate (A3, B0-1, C3) | Severe CAA | 10h | Multiple ICH, CMB |
| 77/M | Alzheimer's disease | Low (A2, B1, C2) | Severe CAA | 11h | Frontal lobe and old temporal ICH |
| 66/F | Alzheimer's disease | Severe (B5, Thal 4, C3) | Mild CAA | 25h | Small subdural |
| 73/M | Alzheimer's disease | Severe (A3, B3, C3) | Moderate CAA | 7h55' | None |
| 82/M | Alzheimer's disease | Low (A3, B0, C2) | Mild CAA | 8h | Hippocampal sclerosis, diffuse Lewey body dementia |
| 65/M | Alzheimer's disease | Severe (A3, B3, C3) | Moderate CAA | 49.5h | Mixed dementia, LBD brainstem |
| 79/M | Alzheimer's disease | Low (A3, B1, C1) | Mild CAA | 7h | None |
| 68/M | Alzheimer's disease | Moderate (A3, B2, C3) | Mild CAA | 18h40' | FTD TDP43, 4R-tau grains |
| 88/F | Alzheimer's disease | Moderate (A3, B2, C2) | Severe CAA | 14h | None |
| 60/F | Alzheimer's disease | Severe (A3, B3, | Mild CAA | 15h | LBD |

| | | | | | |
|---|---|---|---|---|---|
| | | C3) | | | |
| 76/M | Alzheimer's disease | Severe (A3, B3, C3) | Severe CAA | 14h | None |
| 76/M | Alzheimer's disease | Severe (A3, B3, C3) | Moderate CAA | 4h | None |
| 76/M | Alzheimer's disease | Moderate (A3, B2, C2) | Moderate CAA | 21h | ICH |
| 67/F | Alzheimer's disease | Severe (BRAAK VI) | Focally severe | 28h | None |
| 65/F | Alzheimer's disease | Severe (BRAAK VI) | Focally severe | 19h | None |
| 92/F | Alzheimer's disease | Severe (BRAAK VI) | Mild | 16h | None |
| 78/F | Alzheimer's disease | Severe BRAAK V | Mild | 6h | None |
| 62/M | Cerebral amyloid angiopathy | None | Moderate CAA | 35h | Hemorrhages |
| 76/F | Cerebral amyloid angiopathy | Mild (A3, B1, C1) | Severe CAA | 38h | None |
| 68/M | Cerebral amyloid angiopathy | Mild (A3, B1, C1) | Severe CAA | 100h | None |
| 62/M | Neurological control | None | None | 11h3' | TIA |
| 68/M | Neurological control | None | None | 17h18' | None |
| 27/M | Neurological control | None | None | 11h15' | None |
| 45/M | Neurological control | None | None | 13h1' | Cerebrovascular atherosclerosis and thrombosis |
| 87/M | Neurological control | None | None | 12h30' | None |
| 64/M | Neurological control | None | None | 67h17' | Hemorrhages |
| 87/M | Neurological control | None | None | 6h12' | LMCA stroke |
| 66/F | Neurological control | None | None | 33h | None |
| 72/M | Neurological control | None | None | 12h9' | None |
| 68/M | Neurological control | None | None | 24h | |

| | | | | | |
|---|---|---|---|---|---|
| 66/M | Neurological control | None | None | 24h | None |
| 70/F | Neurological control | None | None | 35h15’ | None |
| 82/M | Neurological control | None | None | 164 | ALS (C9orf72) |
| 24/F | Neurological control | None | None | 6h | None |
| 68/F | Neurological control | None | None | 26h | Progressive supranuclear palsy-tau, Parkinsonism |
| 65/F | Neurological control | None | Mild | 17h | Some Lewey body dementia |

SUPPLEMENTARY MATERIAL AND METHODS

**Human brain tissue.**
Brain tissue was obtained from the Vanderbilt Brain and Biospecimen Bank at Vanderbilt University Medical Center, Nashville, Tennessee, USA (IRB# 180287). Written informed consent for each anatomical donation was obtained from patients or their surrogate decision-makers. The Vanderbilt University Medical Center Institutional Review Board provided ethical supervision. The study was conducted in accordance with the ethical principles of the Declaration of Helsinki for experiments involving human subjects.

**Antibodies and reagents.**
*Primary antibodies used in this study include:* rabbit anti-PLD3 (HPA012800, Sigma-Aldrich, St. Louis, MO, 1:1000 for WB and 10 mg/mL for IHC), mouse anti-Ubiquitin clone Ubi-1 (13-1600, Invitrogen, Carlsbad, CA, 1:1000 for WB and 10 mg/mL for IHC), mouse anti-TFEB (91767, Cell Signaling Technology, Boston, MA, 1:1000 for WB, 1:100 for IF), rabbit anti-TFE3 (81744, Cell Signaling Technology, 1:1000 for WB), mouse anti-Lamin B1 (66095, Proteintech, Rosemont, IL, 1:1000 for WB), mouse anti-LC3B (83506, Cell Signaling Technology, 1:1000 for WB), mouse anti-LAMP2 (H4B4, DSHB, Iowa, IA, 1:1000 for WB and 10 mg/mL for IHC), mouse anti-GAPDH (ab59164, Abcam, Cambridge, UK, 1:5000 for WB), mouse anti-GAPDH (60004-1-Ig; Proteintech, 1:1000 for WB), mouse anti-GAPDH-HRP (60004; Proteintech, 1:1000 for WB), rabbit anti-β-Tubulin (15115, Cell Signaling Technology, 1:5000 for WB), goat anti-CTSB (AF965, R&D Systems, Minneapolis, MN, 1:1000 for WB and 10 mg/mL for IHC), goat anti-CTSD (AF1014, R&D systems, 1:1000 for WB and 10 mg/mL for IHC), rabbit anti-IKKβ (8943, Cell Signaling Technology, 1:1000 for WB), normal rabbit IgG (12-370, Cell Signaling Technology).

*Secondary antibodies used in this study include:* goat anti-mouse IgG (H+L), HRP (62-6520; Invitrogen), goat anti-rabbit IgG (H+L), HRP (65-6120; Invitrogen), IRDye® 800 CW goat anti-rabbit (926-32211; LI-COR, Lincoln, NE), IRDye® 680 RD goat anti-mouse (926-68070; LI-COR), IRDye® 680 LT donkey anti-goat (926-68024; LI-COR), IRDye® 800 CW donkey anti-rabbit (926-32213; LI-COR), and IRDye® 800 CW donkey anti-mouse (926-32213; LI-COR).

*Reagents used in this study include:* synthetic β-amyloid 1-42 (AG968; Sigma-Aldrich), dextran Oregon Green 488 (D7172, ThermoFisher Scientific), Protein G/A agarose supernatant (IP05, Millipore, Burlington, MA), dextran Alexa Fluor 594 (D22913, ThermoFisher Scientific), dextran Alexa Fluor 647 (A34785, ThermoFisher Scientific), CellLight Lysosome-RFP BacMam 2.0 (C10597, ThermoFisher Scientific), Torin-1 (14379, Tocris, Minneapolis, MN), Hoechst 33342 (R37605, ThermoFisher Scientific, two drops/ml), neurobasal-A medium, minus phenol red (12349015, ThermoFisher Scientific), 1 µM of 4,4'-[(2-methoxy-1,4-phenylene) di-(1*E*)-2,1-ethenediyl] bisphenol (MX-04) (4920, Tocris), 1 µM Thiazine Red (2150-33-6, Chemsavers Inc, Bluefield, VA), 1 µM of 4 ', 6-diamidino-2-phenylindol (5748, DAPI, Tocris), 1 µM TO-PRO™-3

(T3605, ThermoFisher Scientific). The information for all other reagents is indicated in the corresponding sections.

**Brain tissue preparation, immunostaining, and proximity ligation assay.**

Human brain tissue was obtained at autopsy and prepared as previously described (1). Immunohistochemical labeling was performed as previously described (2-5).

PLD3-ubiquitin complex detection in postmortem human brain was performed using the Duolink® Proximity Ligation Assay (PLA) kit (DUO9200, Sigma-Aldrich), following the protocol previously published (6, 7), including modifications optimized for a fluorescent format in human brain (1). Confocal images were acquired through the Vanderbilt Cell Imaging Shared Resource (CISR) using the Zeiss LSM 710 confocal laser-scanning microscope (Carl Zeiss AG, Germany) with a 20x air/dry or 63x oil objective and 10 µm z-stack scanning projections with a step interval of 1 µm or one scanning projection, with a minimum resolution of 2000 x 2000 pixels. PLA puncta quantification was performed using the HCS Studio software associated with the Cell Insight CX7 high-content imaging system (ThermoFisher Scientific).

**Human β-amyloid seeds extraction.**

Human β-amyloid seeds were isolated as previously described (8) with minor modifications. Seeds were isolated from the brain tissue of patients with severe cerebral amyloid angiopathy (CAA) and minimal other neuropathology. Tissue was homogenized, and the lipid-rich fraction was removed by centrifuging in 17% dextran (40,000 MW, Sigma-Aldrich) at 10,000g for 5 minutes. The pellet was resuspended in 1x TBS, and the suspension was filtered through a custom-designed 3D printed micro sieve apparatus with 40 µm pore size to isolate microvascular fragments. Aggregates of β-amyloid were freed from the vascular matrix by incubating with 3 mg/ml Collagenase Type I (17100017, Gibco, Billings, MT) overnight at 37°C and gently homogenizing with 12 Dounce homogenizer passes. The suspension was again passed through the micro sieve and the liquid containing the particles was centrifuged and resuspended in 1x TBS. It was pelleted and resuspended twice more to remove any residual enzyme thoroughly. The particles were disrupted with sonication to produce a microparticulate suspension with a mean particle diameter of around 10 µm (particle size distribution shown in Fig. S5). Before adding to cells, particles were further disrupted with sonication to produce a microparticulate suspension with a mean particle diameter of about 1 µm (Fig. S5).

**PLD3 knockout cell line.**

The SH-SY5Y-WT human neuroblastoma cell line (CRL-2266, ATCC, Manassas, VA) was used to generate PLD3 knockout (SH-SY5Y PLD3$^{-/-}$), performed by Abcam. PLD3-g1: AGCCCACAACCGCCAGAATG and PLD3-g2: TACTCGCAAGGGTCATAGCA were transfected into SH-SY5Y cells individually with the Cas9 gene to generate the PLD3 knockout two PLD3 knockout clones (C3 and C14). A DNA deletion (~100 bp) in exon 5 of the PLD3 gene in both alleles was achieved, making the frameshift knockout. PCR confirmed the DNA deletion and the knock-out was further validated at the protein level by western blotting.

**Cell culture, cDNA transfection, siRNA, differentiation to neurons, and drug incubation.**
SH-SY5Y-WT and SH-SY5Y PLD3$^{-/-}$ cells were grown in DMEM/F12 (Gibco) supplemented with 10% (v/v) fetal bovine serum (FBS, 35-015-CV, Corning, Corning, NY) and 100 µg/mL Normocin (ant-nnr-1, InvivoGen, San Diego, CA) or 1% penicillin/streptomycin (0503, ScienceCell, Carlsbad, CA) at 37°C in a humidified atmosphere of 5% $CO_2$.
PLD3 plasmids were from Sino Biological (Beijing, China), and the PLD3 mutant plasmids were previously generated using site-directed mutagenesis kit (2). GFP-N-TFEB (38119), and its corresponding control (60360) plasmids were from Addgene (Watertown, MA). PLD3 siRNAs and their corresponding control siRNAs were purchased from Integrated DNA Technologies IDT (Coralville, IA). For plasmid transfection, plasmid DNA was transfected into cells with Lipofectamine 3000 reagent (L3000015, Invitrogen). For siRNA transfection, cells were transfected with 10 mM siRNA by Oligofectamine™ reagent (12252-011, Invitrogen).
SH-SY5Y differentiation into neurons was performed as previously described (9). Cells were differentiated in 10-cm dishes, 8-well chambers slide, 12-well plate or 96-well plate coated with 1 µg/cm$^2$ Matrigel® Matrix (Corning) or poly-L-lysine (P4707, Sigma-Aldrich) and laminin (3400, R&D systems) before performing immunostaining or dextran labeling, protein extraction, SDS-PAGE, and western blot within 10 days of terminal differentiation as described below. Cells were incubated with and without 1 µg/mL of synthetic β-amyloid 1-42 for 1, 3, 5, or 7 days before performing assays. Cells were seeded at a concentration of 1 x $10^5$ per well in pre-coated 8-well chamber slides (154941, ThermoFisher Scientific) and allowed to attach overnight. The cells were then subjected to treatment with DMSO (D2650, Sigma Aldrich), dextran (0.25 mg/ml), HBSS (14025092, Gibco), or Torin 1 (1 µM, Tocris) for three hours at 37°C, followed by washing with 1x PBS for 10 minutes. The cells were then fixed in ethanol containing 5% glacial acetic acid for 12 minutes at -20°C following ICC. Subcellular fluorescence patterns were detected, imaged, and analyzed using Cell Insight CX7 LED Pro HCS Platform.
Differentiation of SH-SY5Y into neuron using retinoic acid was performed as previously described (9)

**Human induced pluripotent stem cell (iPSC) maintenance and differentiation to neurons.**
CC3 iPSCs (10) were maintained and differentiated into cortical glutamatergic neurons as previously described (11) with minor modifications (12). Neurons were used for experiments after at least 70 days of differentiation. iPSCs neurons were plated in an 8-well chamber slide or 12-well plate coated with 1 µg/cm$^2$ Matrigel® Matrix (Corning) before performing immunostaining as described above. iPSC neurons in culture were incubated with and without 100 ng/mL of purified human β-amyloid seeds for 2, 3, 5, or 7 days before performing protein extraction, SDS-PAGE, and western blot as described below, or immunocytochemistry as described above.

**Cathepsin B activity measurement.**
SH-SY5Y-WT and SH-SY5Y PLD3$^{-/-}$ cells were grown in DMEM/F12 (Gibco) supplemented with 10% (v/v) FBS (Corning) and 100 µg/mL Primocin (ANTPM1, InvivoGen) at 37°C in a humidified atmosphere of 5% $CO_2$. Cells were plated in 75 cm$^2$ flasks at a concentration of 5 x $10^6$ cells and cultured overnight before the assay. Cells were collected, and the CTSB enzymatic activity was measured using SensoLyte® 520 Cathepsin B Assay Kit Fluorometric (AS-72164; AnaSpec, Fremont, CA). Results were normalized to the total protein concentration determined by Pierce™ BCA Protein Assay Kit (23225; ThermoFisher Scientific).

**Protein extraction, SDS-PAGE, and Western blot.**
Cell protein homogenates, grey matter, and white matter tissue were separated from human temporal lobe tissue and brain homogenates prepared as described previously (10). Cell protein extraction was performed using RIPA buffer (R0278, Sigma-Aldrich) (10). 20 µg of total protein was resolved in precast 4-20% gradient Mini-PROTEAN® TGX™ gels (Bio-Rad, Hercules, CA) and run at 120V. Proteins were transferred onto Trans-Blot Turbo Mini 0.2 µm PVDF membranes (Bio-Rad) using Trans-Blot® Turbo™ (Bio-Rad). The membranes were dehydrated at 37°C for 10 minutes and soaked with methanol to enhance protein binding. Subsequently, membranes were blocked for 1 hour with 5% BSA (A30075, RPI, Mount Prospect, IL) for chemiluminescence signal development or with Intercept® (TBS) Blocking Buffer (LI-COR) for fluorescent signal development. Primary antibodies were incubated at 4 ˚C overnight and after extensive washes to remove non-specific binding, were incubated with secondary antibodies for 1 hour. Membranes were again washed and developed using SuperSignal™ West Femto Maximum Sensitivity Substrate (34094, ThermoFisher Scientific) and the Chemidoc™ Imaging System (12003154, Bio-Rad).

**Lysosome labeling with dextran.**
For lysosome labeling with dextran, cells were seeded at a concentration of 4 x $10^4$ cells per well in Nunc microwell 96-well optical-bottom plates and allowed to attach overnight. The cells were then incubated with dextran Oregon Green (0.25 mg/ml, D7170, Invitrogen) and dextran Alex Fluor 594 (0.25 mg/ml, D22913, Invitrogen) overnight at 37℃, followed by a chasing period of 6 hours to minimize endosome labeling. The staining medium was removed, and the cells were incubated with Neurobasal medium containing Hoechst for 2 hours at 37℃. Cell Insight CX7 then analyzed the subcellular fluorescence using the LED Pro HCS Platform.

**Co-localization analysis of dextran and LAMP1-RFP.**
Cells were seeded at 4 x $10^4$ cells per well in Nunc microwell 96-well optical-bottom plates and allowed to attach overnight. A construct expressing LAMP1-fused RFP was delivered into cells using BacMam 2.0 technology (C10597, ThermoFisher Scientific) to target lysosomes, following the manufacturer's protocol. On the second day, the medium was removed, and the cells were incubated with dextran Alexa Fluor 647 (0.25mg/ml, A34785, Invitrogen) overnight in the incubator, followed by a chasing period of six hours. The staining medium was removed, and the

cells were stained with Hoechst (Invitrogen) in neurobasal-A medium (Gibco) without dyes for 2 hours at 37°C. CellInsight CX7 then analyzed the subcellular fluorescence using the LED Pro HCS Platform.

**Lysosomal function assay.**
Cells were seeded at 4 x $10^4$ cells per well in Nunc microwell 96-well optical-bottom plates and allowed to attach overnight. The cells were then treated with DMSO (D2650, Sigma Aldrich), Torin-1 (1 μM, Tocris), or Bafilomycin A (100nM, 11038, Cayman Chemical, Ann Arbor, MI) for 3 hours, followed by incubation with DQ Green BSA (10 ug/ml, D12050, ThermoFisher Scientific) and Alexa Fluor 647 BSA (10 ug/ml, A34785, ThermoFisher Scientific) for 3 hours in the incubator. The assay medium was removed, and the cells were incubated with Hoechst in neurobasal-A medium for nuclear staining for 2 hours at 37°C. Subcellular fluorescence was analyzed using the Cell Insight CX7 LED Pro HCS Platform.

**Spot detection and quantification by CellInsight CX7 LED Pro HCS platform.**
Subcellular fluorescence was detected and imaged in acquisition mode on the CX7 HCS platform. Images *(n=30-50)* per well were automatically captured using laser autofocus, and the in-focus pictures were selected for subsequent analysis. For quantification, images were first preprocessed with background removal, smoothing, and segmentation, followed by spot (fluorescent signals) identification and validation. The spots were compartmentalized into the cell body (circle) and the distal part (ring). All images were re-scanned in disk mode, and the information about spot area and intensity (total and average) on all channels was statistically analyzed. Three to six independent experiments (n=3-6) were performed for final statistical analysis.

**Transmission electron microscopy.**
Cells were fixed in 0.1 M cacodylate buffer containing 2.5% glutaraldehyde, harvested using a cell scraper, and pelleted. Samples were then sequentially postfixed in 1% tannic acid and 1% OsO4, followed by en block staining in 2% uranyl acetate. After post-fixation, the samples were dehydrated in a graded ethanol series and infiltrated with Quetol 651-based Spurr's resin using propylene oxide as a transition solvent. The resin was polymerized at 60 °C for 48 hours, and 70 nm sections were cut on a Leica UC7 ultramicrotome.
Imaging was performed on a JEOL 2100+ transmission electron microscope operating at 200 keV, with images captured using an AMT nanosprint CMOS camera. Single images were acquired with AMT imaging software, while tiled imaged sets were acquired using SerialEM automation software. Tiled datasets were processed using the IMOD/etomo software suite and FIJI. Lysosome characteristics were analyzed and quantified using FIJI.

**Lysosomal protein preparation.**
Lysosomes were isolated and purified as described previously (2). Briefly, cells were grown to around 80% confluency and incubated with DMEM/F12 (Gibco) medium containing 10 mg/ml

dextran-coated iron oxide nanoparticles for 8 hours. The cells were washed three times with complete DMEM/F12 medium and rested in an incubator for 16 hours. The cells were collected in homogenization buffer (containing protease inhibitors) and homogenized with a Dounce homogenizer for 12 strokes. The lysates were centrifuged, and the supernatants, post-nuclear supernatant (PNS), were then passed through a pre-equilibrated column that was pre-filled with fine steel fibers and embedded in a magnet. The flow-through (FT) was collected after being applied to the column three times. Next, the column was washed with homogenization buffer three times and removed from the magnet. The lysosome fraction (LF) in the column was then eluted and lysed with buffers that are compatible with the following experiments. For western blot analysis, RIPA buffer (Sigma-Aldrich) was used; for proteomic study, PBS containing 1% SDS was used.

**Immunoprecipitation (IP).**

Cells were washed with PBS 1x and then lysed in Pierce IP lysis buffer (87787, ThermoFisher Scientific). Approximately 500 µg of lysates was pre-cleared with 0.5 µg of normal rabbit IgG and 10 µl of protein G/A agarose solution, followed by incubation for 1 hour at room temperature. The supernatant was subsequently incubated overnight with 30 µL of protein G/A agarose solution and 5 µg of TFE3 antibody or rabbit normal IgG (IP control). On the following day, agarose beads were briefly centrifuged and washed three times with PBS. The agarose pellets were then reconstituted with 40 µl of 2x Laemmli protein buffer (1610747, Bio-Rad), followed by Western blot analysis to detect ubiquitinated TFE3.

**Isolation of cytoplasmic and nuclear proteins.**

Cytoplasmic and nuclear proteins were isolated from cells of interest following the instructions in the manufacturer's user guide (78833, ThermoFisher Scientific). Briefly, cells were detached with trypsin-EDTA treatment and pelleted by centrifuge at 500 g x for 5 minutes. The cells were then washed once with 1x PBS. After the 1x PBS wash, the cell pellet was resuspended in CER I buffer, vortexed and incubated on ice for 10 minutes. CER II buffer was then added to the suspension and mixed, followed by incubation for 1 minute. The suspension was then centrifuged, and the supernatant was immediately transferred to another tube (cytoplasmic proteins). The remaining nuclear pellet was resuspended in ice-cold NER buffer and incubated on ice for at least 40 minutes with intermittent vortex every 10 minutes. The suspension was then centrifuged, and the supernatant was collected in a new tube (nuclear proteins). The volume ratio of CER I: CER II: NER reagents was maintained and proportional to cell numbers as instructed in the user guide.

**Mass spectrometry and bioinformatic analysis.**

Protein samples were prepared with 5% SDS, processed using S-Trap (ProtiFi) digestion, and analyzed by LC-MS/MS, following a protocol described previously (13). Proteins were first reduced with 10 mM TCEP, alkylated with 20 mM iodoacetamide, and then 2.5% aqueous phosphoric acid was added, which is followed by the addition of S-Trap binding buffer (90% methanol in 100 mM TEAB) at six times the sample volume. The samples were loaded onto S-Trap microcolumns, washed with binding buffer, and digested with trypsin gold (Promega, Madison, WI) at a 1:10 enzyme-to-protein ratio in 50 mM TEAB (pH 8.0) for 1 hour at 47°C. Peptides were eluted in series with 40 µL each of 50 mM TEAB, 0.2% formic acid, and 35 µL of 0.2% formic acid in 50% acetonitrile. The eluted peptides were dried and reconstituted in 0.1% formic acid for LC-MS/MS analysis. Peptides were loaded onto a C18 reverse-phase analytical column using a Dionex Ultimate 3000 nanoLC system with autosampler and eluted using a 90-minute gradient (1–72 min: 2–38% B; 72–78 min: 38–90% B; 78–80 min: 90% B; 80–81 min: 90–2% B; 81–90 min: 2% B for re-equilibration). The Orbitrap Exploris 240 mass spectrometer (Thermo Scientific), equipped with a nanoelectrospray ionization source, was used for data-dependent acquisition. The instrument method included MS1 scans with an MS AGC target of $3 \times 10^6$, followed by 20 MS/MS scans of the most abundant ions detected in each MS1 scan. The intensity threshold for triggering MS/MS scans was set at $1 \times 10^4$, with an MS2 AGC target of $1 \times 10^5$. Dynamic exclusion was set to 10 seconds, and HCD collision energy was 30 nce. The LC-MS/MS data were analyzed using MaxQuant against a human database created from the UniprotKB protein database, with MaxQuant contaminants added (14). Standard Maxquant parameters were applied, along with LFQ and match-between-runs. Variable modifications included methionine oxidation and N-terminal acetylation, while carbamidomethyl cysteine was set as a fixed modification. A false discovery rate (FDR) of 0.01 was applied to peptide and protein identifications. Label-free quantitative (LFQ) analysis of the identified proteins was performed using the MSstats R package with default parameters (15).

**RNA extraction, sequencing, and bioinformatic analysis.**

Total RNA was isolated from cells of interest using RNeasy Mini Plus Kits (74134, Qiagen, Netherlands) following the manufacturer's protocol, with each group having three replicates. Two micrograms of RNA per sample were submitted to Vanderbilt Technologies for Advanced Genomics (VANTAGE) for quality control analysis, library preparation, and next-generation sequencing (NGS). RNA quality control was performed using the Agilent Bioanalyzer and RNA quantity was determined using the RNA Qubit assay (Q10210, ThermoFisher Scientific).

**Statistical analysis**
The number of samples (*n*) in each group/experimental condition is indicated in the figure legends. All data were graphed and statistically analyzed using GraphPad software (version 10.0.2, GraphPad Software Inc., San Diego, CA) or R packages unless stated otherwise. For gene set enrichment analysis, statistical tests were based on the Kolmogorov-Smirnov test (18). For RNA-Seq differential gene expression analysis, the data were subject to negative binomial generalized linear model fitting and Wald statistics (17). To compare between two groups, parametric unpaired *t*-test or Welch's unequal variances *t*-test were used. To compare more than two independent groups, ordinary one-way ANOVA was used followed by Tukey’s or Dunnett multiple comparisons, or non-parametric Kruskal-Wallis tests were performed followed by Dunn’s multiple comparisons test. To assess the individual and interaction effects of two variables, two-way ANOVA tests were performed, followed by the Dunnett test. Statistical tests were specified and noted in the respective figure legends. A significant level of 0.05 was used to reject the null hypothesis throughout this article unless noted otherwise.

### SUPPLEMENTARY FIGURE LEGEND

**Fig. S1. Validation of PLD3 antibody by western blot and immunostaining.**
(A) PLD3 western blot band pattern in wild type SH-SY5Y (WT) cells and in cells with genetically knocked out PLD3 ($PLD3^{-/-}$) are shown.
(B) Representative confocal microscopy images of PLD3 immunostaining *(green)* in the SH-SY5Y-WT cells and SH-SY5Y-$PLD3^{-/-}$ cells. Nuclei were stained with DAPI *(blue).* The scale bars are indicated.

**Fig. S2. PLD3 is reduced in human hippocampus, posterior frontal cortex and superior temporal gyrus in AD.**
(A-B) Representative confocal microscopy images of PLD3 immunostaining *(green)* in the hippocampus from non-AD (A) and AD brain (B) sections, *n=8.* Nuclei were stained with DAPI *(blue),* and β-amyloid and neuritic plaques, neurofibrillary tangles, and other tau aggregates were stained with Thiazine Red *(TR, red)*. The scale bars are indicated.
(C-D) Representative confocal microscopy images of PLD3 immunostaining *(green)* in posterior frontal (PF.C) (C), and superior temporal gyrus (T.C) (D) sections from non-AD *(left panel)* and AD *(right panel)* brain sections, *n=3.* β-amyloid and neuritic plaques, neurofibrillary tangles, and other tau aggregates were stained with Methoxy-X04 *(MX-04, blue)* or Thiazine Red *(TR, red)*. Nuclei were stained with DAPI *(blue).* The scale bars are indicated.

**Fig. S3. PLD3 locate with neuronal but not with astrocyte marker in human brain.**
(A) Representative confocal microscopy images of PLD3 immunostaining *(green)* in grey matter vs. white matter, *n=3.* Nuclei were stained with DAPI *(blue).* The scale bars are 1000 µm.
(B) 299 individuals with normal cognition control (*n= 142*, cogdx = 1) or AD dementia (*n = 157*, cogdx = 4 or 5) from the ROSMAP cohort were analyzed for differential gene expression across 8 cell types, demonstrating the PDL3 is primarily expressed in neurons.
(C-E) Representative confocal microscopy images of colocalization of the PLD3 immunostaining *(green)* with neuron marker (Neurofilament, NF) (C), microglia marker (Ionized Calcium-Binding Adaptor Molecule 1, Iba1) (D) and astrocyte marker (Glial Fibrillary Acidic Protein, GFAP) (E) in grey matter *(left panel)* and white matter *(right panel)* from non-AD brain sections, *n=3.* The scale bars are 50 µm.
(F) Representative confocal microscopy images of colocalization of the PLD3 immunostaining *(green)* with microglia marker (Ionized Calcium-Binding Adaptor Molecule 1, Iba1) in non-AD *(left panel)* and AD *(right panel)* brain sections, *n=3.* The scale bars are 50 µm.
(G) Strength of the linear relationship between PLD3 with Iba1 immunostaining. Pearson correlation coefficient *(PCC, (r))* is presented as mean ± S.E.M., *n=6.*

**Fig. S4. Raw data for PLD3 levels in grey matter and white matter.**

(A) Western blot of PLD3 and GAPDH in superior temporal gyrus protein lysates from grey matter. Each lane corresponds to a donor of the indicated group.
(B) Western blot of PLD3 and GAPDH in superior temporal gyrus protein lysates from white matter. Each lane corresponds to a donor of the indicated group.

**Fig. S5. Influence of brain-derived β-amyloid seeds and synthetic Aβ$_{1-42}$ on PLD3 expression.**
(A) Raw images of western blots assessing effect of 7-day incubation with synthetic Aβ$_{1-42}$ (1 μg/mL) on PLD3 and GAPDH levels in SH-SY5Y-WT cells.
(B) Analysis of the size of human amyloid seeds isolated from human brain tissue with cerebral amyloid angiopathy stained with Thiazine Red *(TR)* using Countess 3 FL (Invitrogen). Particles before adding to cells were further disrupted with sonication to produce a microparticulate suspension with a mean particle diameter of about 1 micron.
(C) Western blot analysis of brain-derived amyloid seeds for β-amyloid (APP Antibody, 6E10, NBP2-62566, Novus Biologicals) and phospho-Tau Ser202-The205 (MN1020, Invitrogen).
(D) Representative confocal microscopy images of β-amyloid seeds immunostaining *(MX-04, blue)* in iPSC-derived neurons. Neuronal cell bodies were stained with Neurofilament *(NF, red)*. The scale bars are 50 μm.
(E) Raw images of western blots assessing the effect of 7-days incubation with brain-derived β-amyloid seeds on PLD3 levels in SH-SY5Y-WT cells.

**Fig. S6. Neuronal lysosome function after exposure to brain-derived β-amyloid seeds.**
(A) Representative confocal microscopy images of SH-SY5Y-WT-derived neurons treated with brain-derived β-amyloid seeds for 1, 3 and 5 days. Lysosomes were labeled with Dextran Oregon Green *(green)*, brain-derived β-amyloid seeds were detected with Dextran Alexa 594 *(red)*, nuclei were stained with Hoechst *(blue)* and cell bodies are shown in brightfield.
(B) Quantification, and comparison of lysosome features (size, positioning, number, total area, and alkalinization) among experimental conditions ($n= 5\text{-}6$, mean ± S.E.M., one-way ANOVA followed by Dunnett's multiple comparison test). The scale bars are 500 nm. NS: non-stimulation.

**Fig. S7. Differential ubiquitin and PLD3 immunostaining in human brain tissue.**
(A) Representative confocal microscopy images of ubiquitin immunostaining *(green)* in white and grey matter for non-AD *(top panel)* and AD *(bottom panel)* human brain. Nuclei were stained with DAPI *(blue)*. β-amyloid, neuritic plaques, neurofibrillary tangles, and other tau aggregates were stained with Thiazine Red *(TR, red)*. The scale bars are indicated.
(B) Representative confocal microscopy image of an *in-situ* proximity ligation assay (PLA) detecting PLD3-ubiquitin complexes *(red)* in AD human brain associated with tau aggregates. Neurofibrillary tangles and other tau aggregates were stained with Methoxy-X04 *(MX-04, blue)*.

(C) Representative confocal microscopy images of PLA validation probes. PLA signal *(red)* using only the anti-PLD3 antibody *(left panel)*, only the anti-ubiquitin antibody *(middle panel)* and just PLA probes *(right panel)*. Nuclei were stained with TO-PRO$^{TM3}$ *(blue)*. β-amyloid, neuritic plaques, neurofibrillary tangles, and other tau aggregates were stained with Methoxy-X04 *(MX-04, green)*. The scale bars are 50 μm.

**Fig. S8. Lysosomal morphology and function in SH-SY5Y-PLD3$^{-/-}$ cells.**
(A) Quantification of relative lysosome: area, size and number per cell. Data are presented as means ± S.E.M. $^{*}p < 0.05$ by Welch's unequal variances *t*-test.
(B) Dextran (green) and Lyso-RFP (red) were analyzed in WT and PLD3$^{-/-}$ cells transduced with Lyso-RFP viruses for 24 hours and then stained with dextran Alexa 647 overnight, followed by a chasing period of at least 6 hours and Hoechst nuclear counterstaining for 2 hours. Scale bars, 10 μm. Colocalization/correlation of dextran and lyso-RFP was assessed by Pearson coefficient and Mander's coefficient and compared between WT and PLD3$^{-/-}$ cells (*n=3*, unpaired t test).

**Fig. S9. Transcriptomic and proteomic analysis of lysosomal dysregulation after PLD3 deletion clone 3 and clone 14.**
(A) GSEA plots featuring upregulated endosomal and lysosomal pathways in SH-SY5Y-WT cells (*WT)* and SH-SY5Y-PLD3$^{-/-}$ cells (*PLD3$^{-/-}$, clone 3*) (FDR q-value < 0.05; nominal p-value <0.05).
(B) Proteomic analysis of the lysosomal lysates from SH-SY5Y-WT cells (*WT)* and SH-SH5Y-PLD3$^{-/-}$ cells (*PLD3$^{-/-}$, KO clone 14*). The differentially expressed proteins were represented by heatmap (n=3, ≥1.5-fold change, false discover rate (FDR) < 0.05). Proteomic data (normalized) were subject to GSEA analysis, and the differentially regulated pathways in in SH-SY5Y-WT cells (*WT)* and SH-SY5Y-PLD3$^{-/-}$ cells (*PLD3$^{-/-}$, KO clone 14*) were summarized and represented by bar plots (FDR q value < 0.05).
(C) GSEA plots featuring the enriched lysosomal pathway in SH-SY5Y-WT cells (*WT)* and SH-SY5Y-PLD3$^{-/-}$ cells (*PLD3$^{-/-}$, clone 14*) (FDR q-value < 0.05; nominal p-value <0.05).
(D) Lysosomal proteins from proteomic analysis were quantified and compared between SY-SH5Y-WT cells (*WT)* and SH-SY5Y-PLD3$^{-/-}$ cells (*PLD3$^{-/-}$, clone 14*), and the differentially regulated proteins were represented by bar graphs (mean ± S.E.M., multiple unpaired t test).

**Fig. S10. Transcriptomic profiling following PLD3 knockout in SH-SY5Y cells.**
(A) RNA sequencing analysis of total RNA from SH-SY5Y-WT cells (*WT)* and SH-SY5Y-PLD3$^{-/-}$ cells (*PLD3$^{-/-}$, KO*). RNA count data were log-transformed (loge) and analyzed using principal component analysis (PCA).
(B) The differentially expressed genes in SH-SY5Y-WT cells (*WT)* and SH-SY5Y-PLD3$^{-/-}$ cells (*PLD3$^{-/-}$, KO*) were represented by heatmap (*n=3*, false discover rate (FDR) < 0.05).

(C) The differentially expressed genes were represented by volcano plots with the differentially expressed lysosomal proteins labeled.
(D) RNA-Seq data were subject to GSEA analysis, and the differentially enriched pathways were represented by bar plots.
(E) GSEA plots featuring the upregulated lysosomal pathway and AD in SH-SY5Y-WT cells (*WT)* and SH-SY5Y-PLD3$^{-/-}$ cells (*PLD3*$^{-/-}$*, KO*).
(F) RNA expression in genes related to the lysosome pathway was calculated and compared between SH-SY5Y-PLD3$^{-/-}$ cells (*PLD3*$^{-/-}$*, KO*) *(n=3)* and SH-SY5Y-WT cells (*WT) (n=3)*, and significantly different genes were represented by bar graphs (mean ± S.E.M., multiple unpaired t test).

**Fig. S11. Transcriptomic profiling after PLD3 knockdown using siRNA.**
(A) Western blot analysis of PLD3 in SH-SY5Y cells transfected with CTL_siRNA or PLD3_siRNAs for 2 days. Proteins were quantified, expressed relative to GAPDH, and compared (*n=3-4*, mean ± S.E.M., one-way ANOVA followed by Dunnett's multiple comparison test).
(B) RNA sequencing analysis of the total RNA in SH-SY5Y cells transfected with CTL_siRNA or PLD3_siRNAs for 2 days. The differentially expressed genes were represented by heatmap (n=3, ≥1.5-fold change, FDR < 0.05).
(C) The differentially expressed genes in cells transfected with PLD3 siRNAs versus CTL siRNA were represented by volcano plots with lysosomal proteins labeled.
(D) RNA sequencing count data were log-transformed (log) and analyzed using principal component analysis (PCA).

**Fig. S12. PLD3 knockdown causes enrichment in lysosome-related signaling.**
(A) GSEA plots featuring upregulated endosomal, autophagic and lysosomal pathways in PLD3_siRNAs versus CTRL_siRNA (FDR q-value < 0.15; nominal p-value <0.05).

**Fig. S13. CTSB expression in Alzheimer's disease human brain.** Western blot of CTSB, GAPDH and NeuN in superior temporal gyrus lysate derived from grey matter from non-AD and AD human brains. Each lane corresponds to a donor of the indicated group.

**Fig. S14. CTSB locate in extra-lysosomal compartments in the human AD brain.**
(A-B) Colocalization of CTSB and LAMP2 in huma brain. Representative confocal microscopy images showing colocalization of CTSB and LAMP2 in non-AD (A) and AD (B) human brains in cortex and hippocampus. β-amyloid, neuritic plaques, neurofibrillary tangles, and other tau aggregates were stained with Methoxy-X04 *(MX-04, blue).* Scale bars are indicated.
(C) CTSB reduce its colocalization with PLD3 in AD brains. Representative confocal microscopy images of colocalization of CTSB and PLD3. β-amyloid, neuritic plaques,

neurofibrillary tangles, and other tau aggregates were stained with Methoxy-X04 *(MX-04, blue)*. Scale bars are indicated.
(D) CTSB is associated with the human brain microvasculature in AD brains. Representative confocal microscopy images of colocalization of CTSB and collagen-IV. β-amyloid, neuritic plaques, neurofibrillary tangles, and other tau aggregates were stained with Methoxy-X04 *(MX-04, blue).* Scale bars are indicated.

**Fig. S15. PLD3 modulates TFEB/TFE3 metabolism without affecting their transcription.**
(A) RNA levels of TFEB and TFE3 from RNA sequencing of total RNA in SHY-SY5Y-WT cells (*WT)* and SH-SY5Y-PLD3$^{-/-}$ cells (*PLD3$^{-/-}$, KO*), or CTL_siRNA and PLD3_siRNA cells, were quantified and expressed relative to the corresponding controls (*n=3*, mean ± S.E.M., t-test).
(B) Western blot analysis of TFEB, TFE3 and PLD3 in SH-SY5Y-WT cells (*WT)* and SHY-SY5Y-PLD3$^{-/-}$ cells (PLD3$^{-/-}$) treated with vehicle (NS), dextran (0.25 mg/ml), HBSS, or Torin 1 (1 μM) for 8 hours. NS: non-stimulation. Proteins were quantified, expressed relative to β-tubulin, and compared between genotypes (*n=3-4*, mean ± S.E.M., two-way ANOVA followed by multiple comparison test).
(C) Western blot analysis of TFEB, TFE3 and PLD3 in SH-SY5Y-WT cells (*WT)* transfected with control siRNAs (CTL_siRNA) or PLD3 siRNAs for 2 days. Proteins were quantified, expressed relative to β-tubulin, and compared between genotypes (*n=3-4*, mean ± S.E.M., one-way ANOVA followed by Dunnett's multiple comparison test).
(D) Western blot analysis of p-TFEB (Ser 211) and p-TFEB (Ser 142) in SH-SY5Y-WT cells (*WT)* and SH-SY5Y-PLD3$^{-/-}$ cells treated with vehicle (NS), dextran (0.25mg/ml), HBSS, or Torin 1 (1 μM) for 8 hours.

**Fig. S16. PLD3 mediates TFEB/TFE3 degradation through the proteasome.**
(A-B) Western blot analysis of TFEB, TFE3 and PLD3 in SH-SY5Y-WT cells (*WT)* and SH-SY5Y-PLD3$^{-/-}$ cells treated with bortezomib (BZ, 200 nM/DMSO) for 4 hours (A) and 8 hours (B). Proteins were quantified, expressed relative to β-tubulin, and compared among experimental conditions (*n=3-4*, mean ± S.E.M., two-way ANOVA followed by Dunnett's multiple comparison test).
(C) Western blot analysis of IKKβ in SH-SY5Y-WT cells (*WT)* and SH-SY5Y-PLD3$^{-/-}$ cells treated with BZ (200 nM) or MG132 (1 μM) for 4 hours. Proteins were quantified, expressed relative to β-tubulin, and compared among experimental conditions (*n=3-4*, mean ± S.E.M., two-way ANOVA followed by Tukey test).
(D) Western blot analysis of p-IKK /β (Ser 176/180) in SH-SY5Y-WT cells (*WT)* and SH-SY5Y-PLD3$^{-/-}$ cells treated with DMSO, BZ, and MG132 for 8 hours.

**Fig. S17. The proteasome, rather than lysosomes, are primarily involved in PLD3-mediated TFEB/TFE3 metabolism.**

(A) Western blot analysis of TFEB, TFE3 and PLD3 in SY-SY5Y-WT cells and SY-SY5Y-PLD3$^{-/-}$ cells treated with cycloheximide (CHX, 20 µg/mL) and/or MG132 (1 µM) for 4 hours (representative images of two independent experiments).
(B) Western blot analysis of TFE3 immunoprecipitation for TFE3 and ubiquitin from the lysates of SY-SY5Y-WT cells and SY-SY5Y-PLD3$^{-/-}$ cells treated with or without MG132 (1 µM) for 4 hours. An immunoprecipitation control with rabbit normal IgG was included to demonstrate the specificity of the immunoprecipitation (representative images of two independent experiments).

**Fig. S18. Loss of PLD3 augments the level of nuclear TFEB/TFE3 and their transcriptional activities.**
(A) Representative confocal microscopy images of immunostaining of TFEB *(green)* in SH-SY5Y-WT cells (*WT)* and SH-SY5Y-PLD3$^{-/-}$ cells treated with HBSS (vs NS) or 1 µM Torin 1 (vs. DMSO) for 3 hours with DAPI nuclear counterstaining *(blue)*. The scale bars are 50 µm. NS: non-stimulation.

**Fig. S1.** Romero-Fernandez W. and Wang Y. *et al.*, 2025.

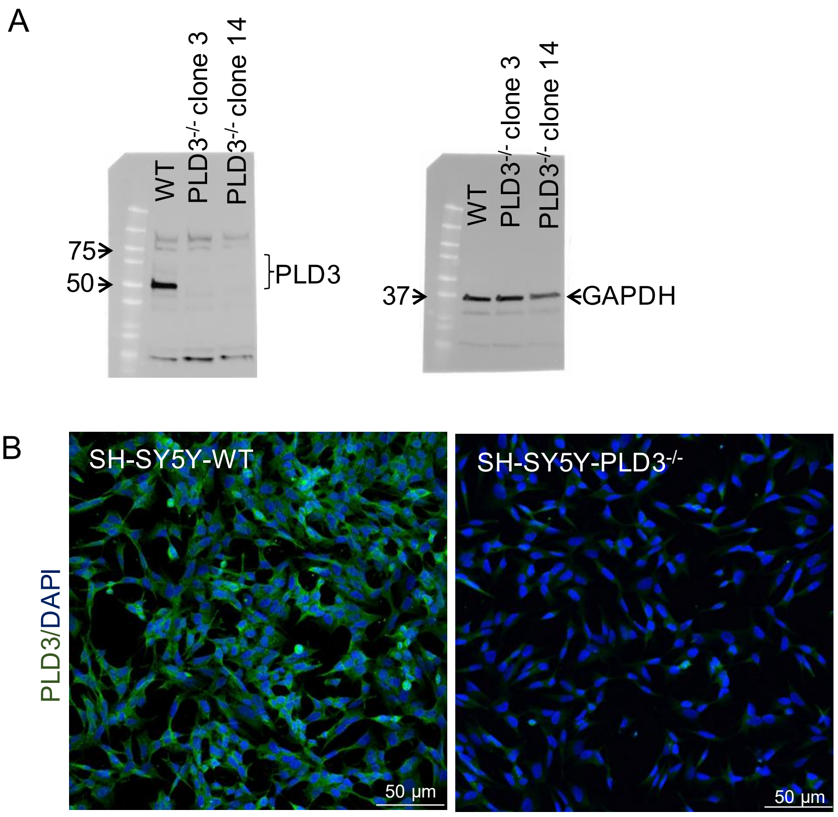

**Fig. S2.** Romero-Fernandez W. and Wang Y. *et al*., 2025.

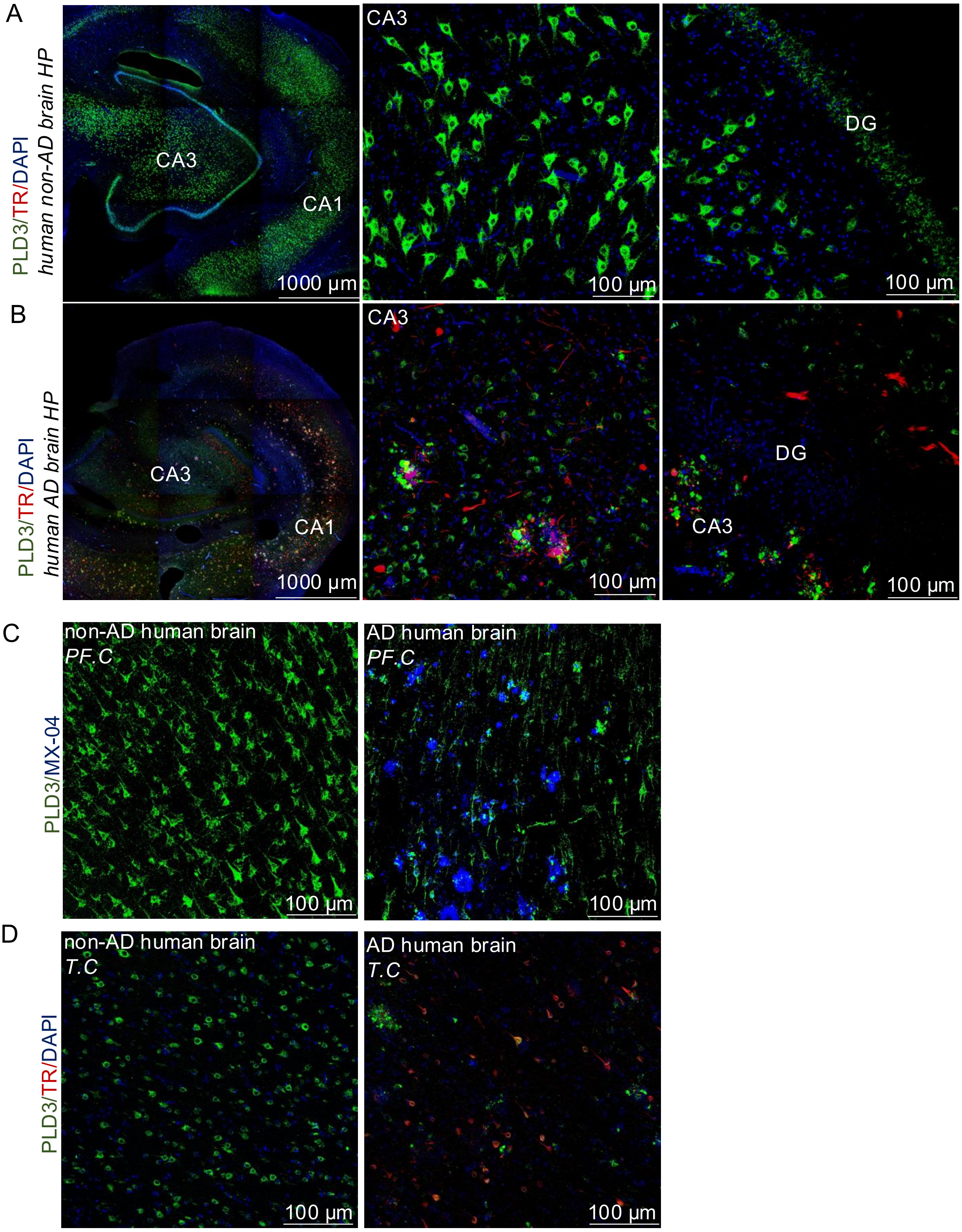

**Fig. S3.** Romero-Fernandez W. and Wang Y. *et al.*, 2025.

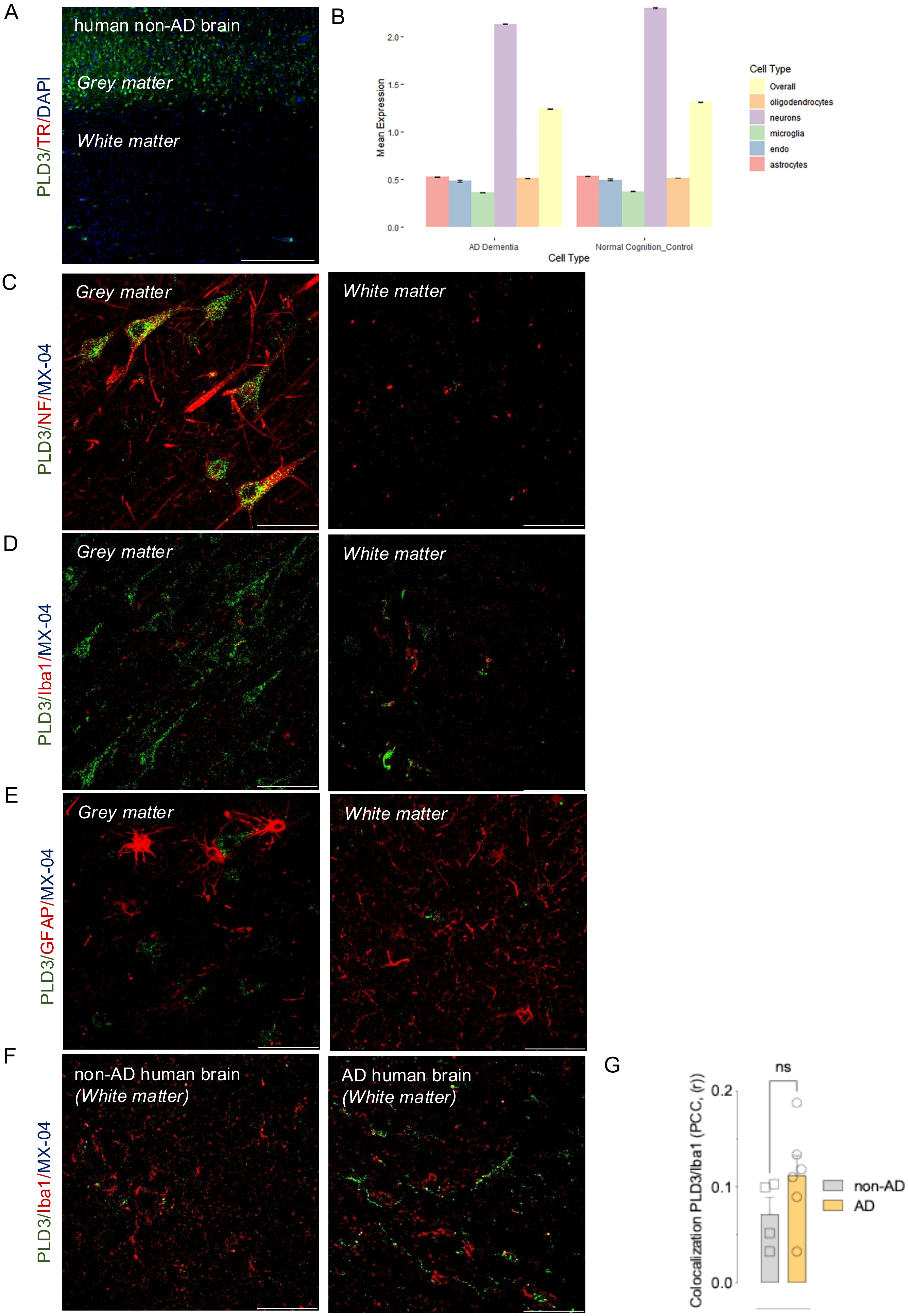

**Fig. S4.** Romero-Fernandez W. and Wang Y. *et al.*, 2025.

A

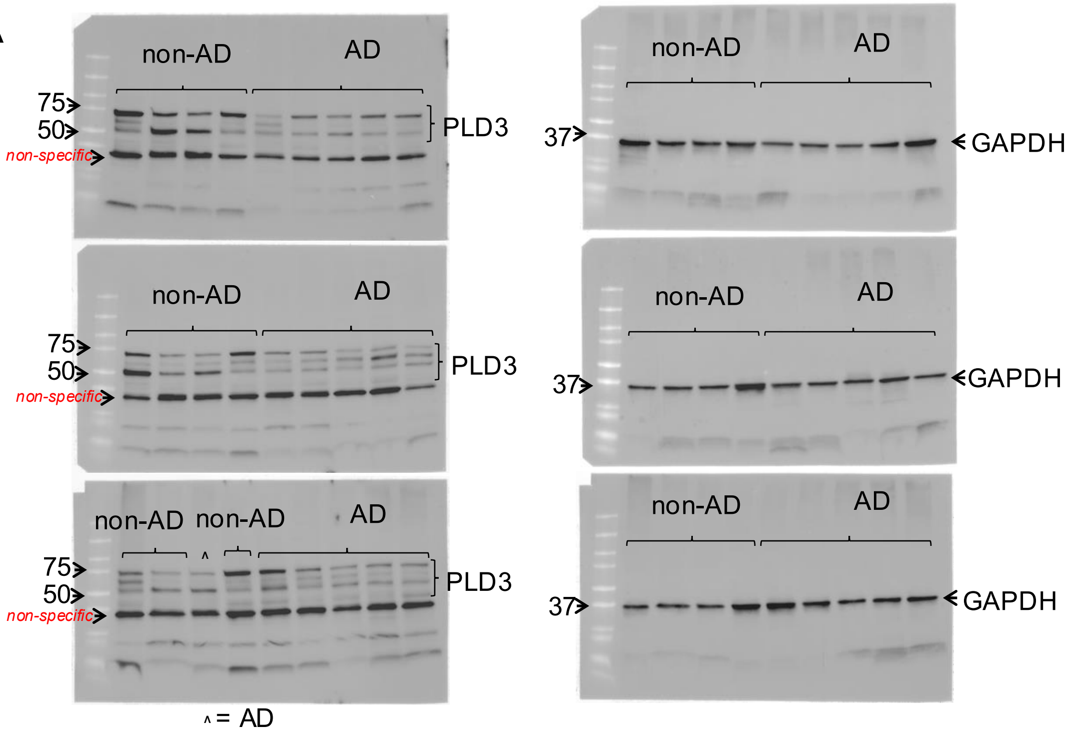


B

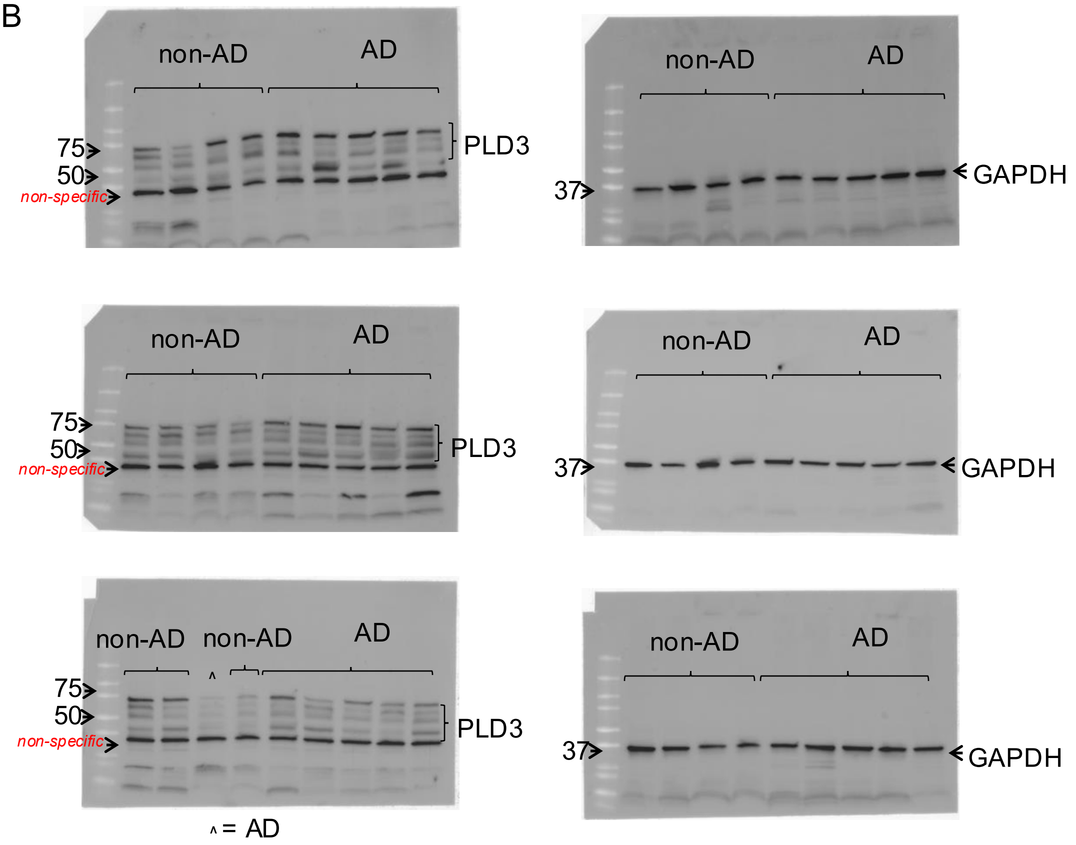

**Fig. S5.** Romero-Fernandez W. and Wang Y. *et al.*, 2025.

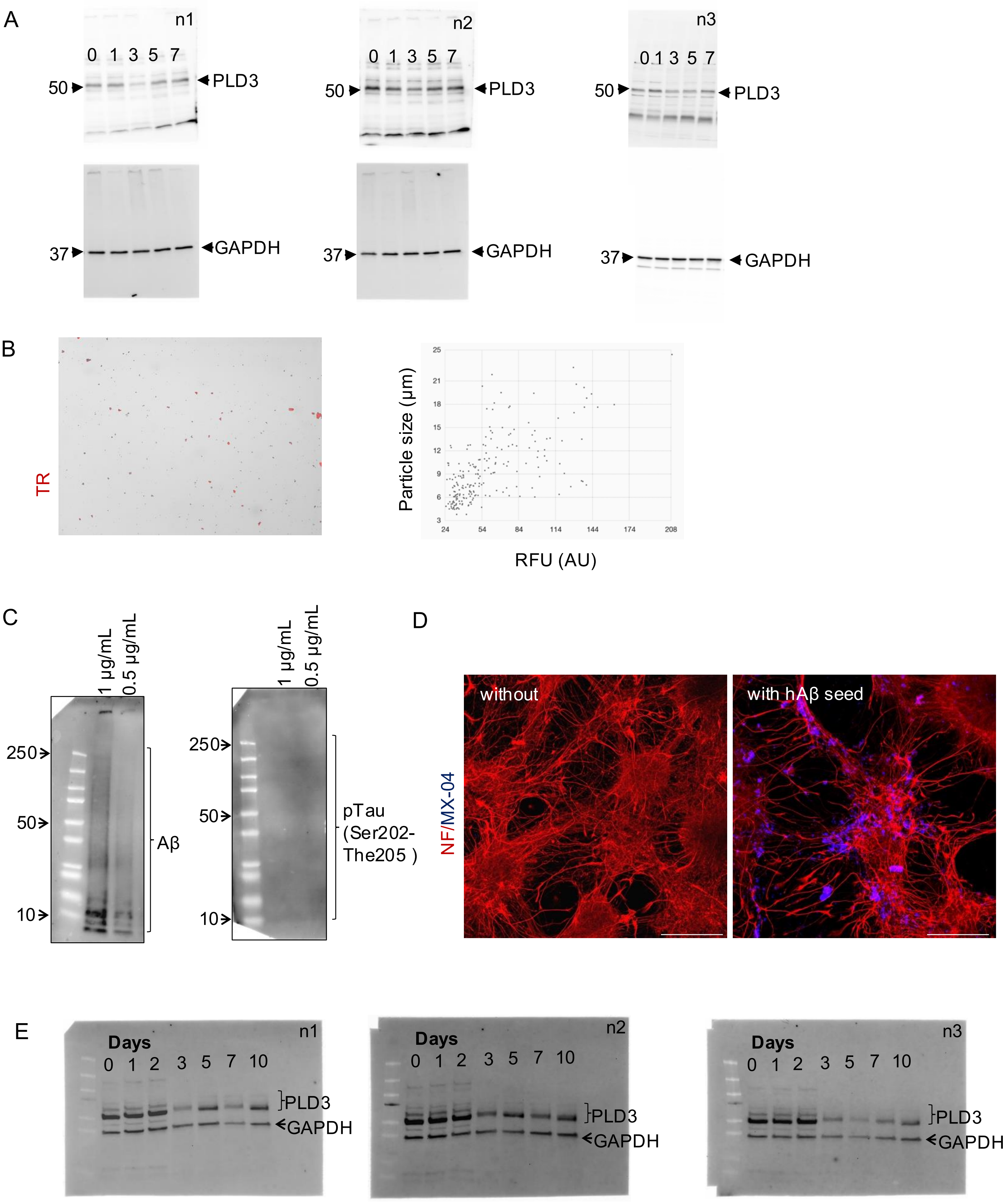

**Fig. S6.** Romero-Fernandez W. and Wang Y. *et al.*, 2025.

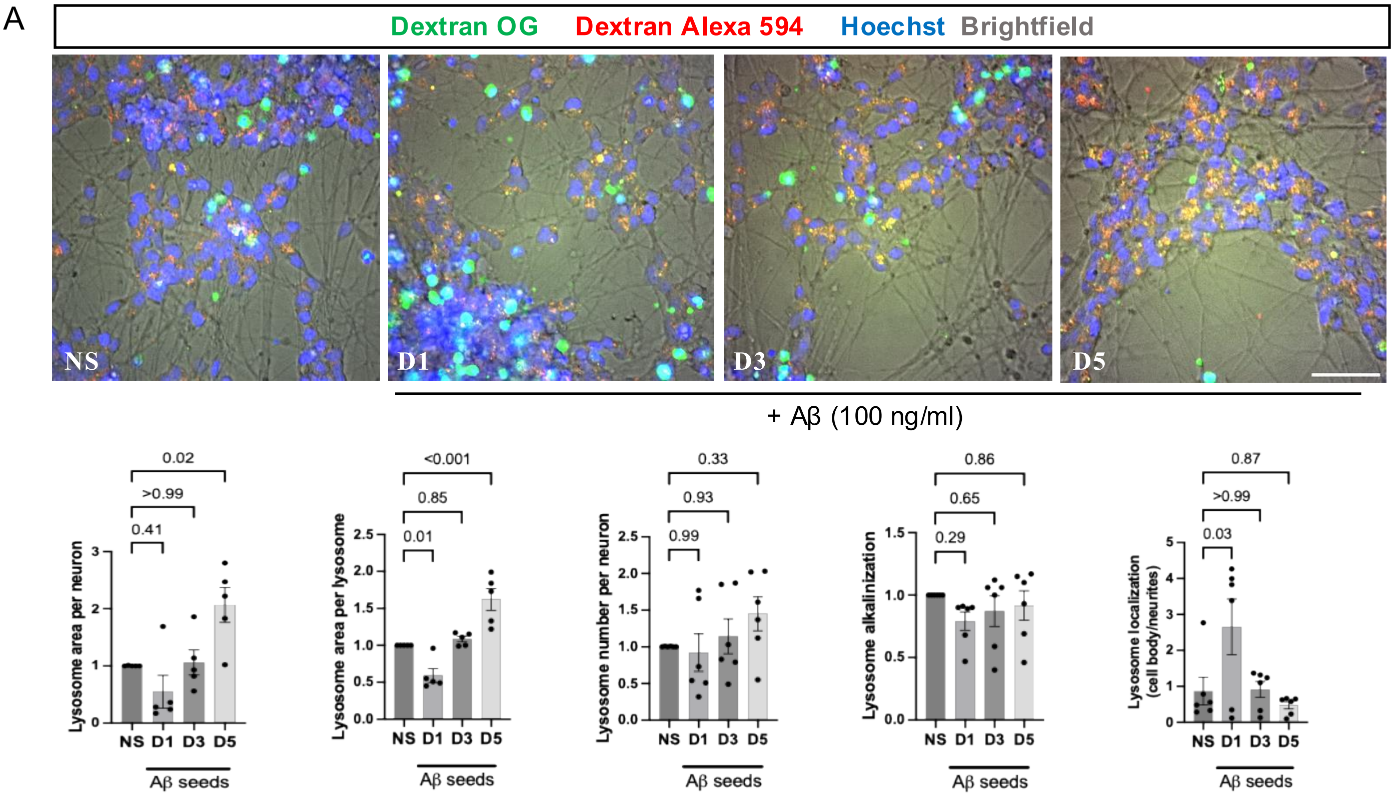

**Fig. S7.** Romero-Fernandez W. and Wang Y. *et al.*, 2025.

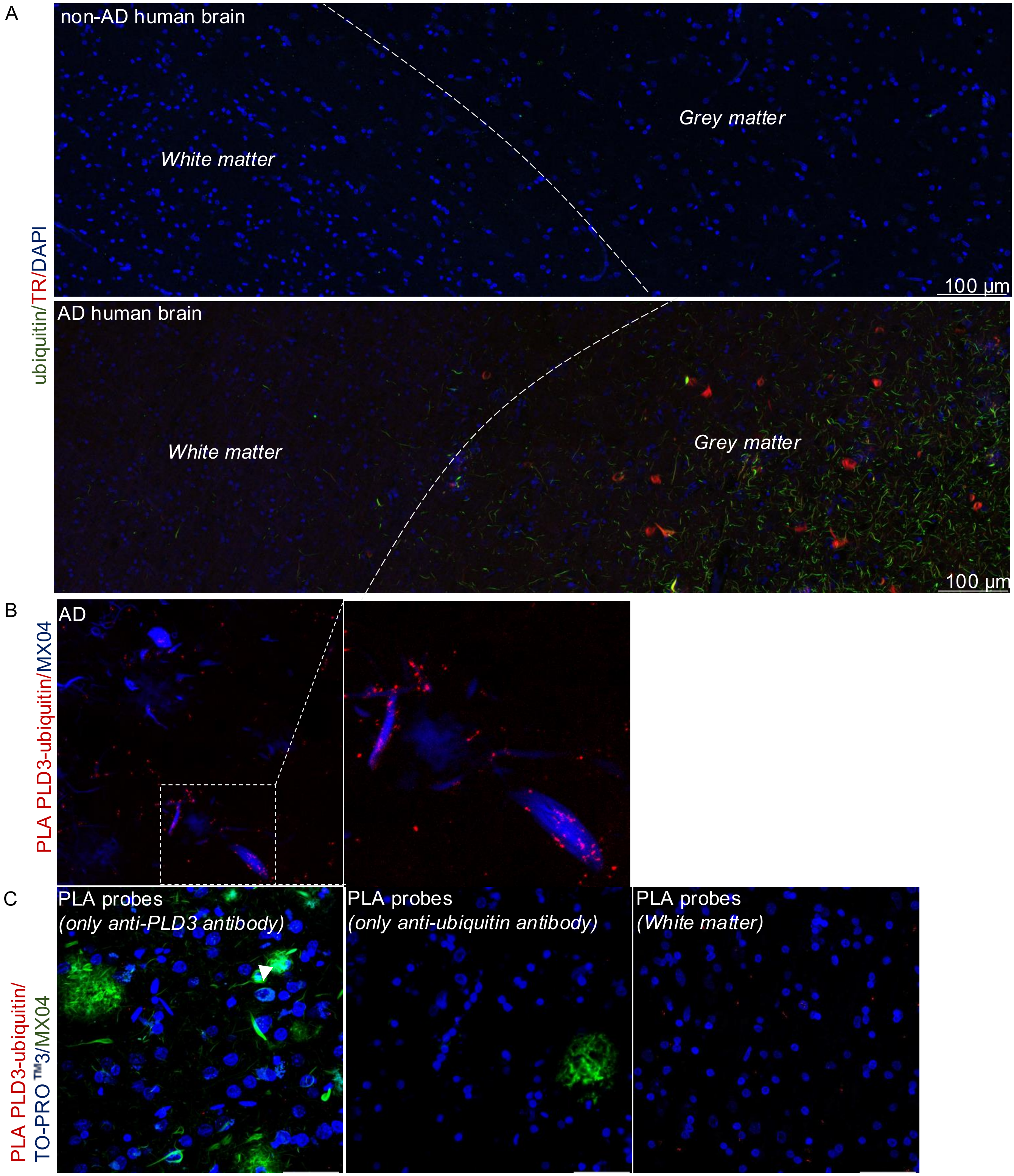

**Fig. S8.** Romero-Fernandez W. and Wang Y. *et al.*, 2025.

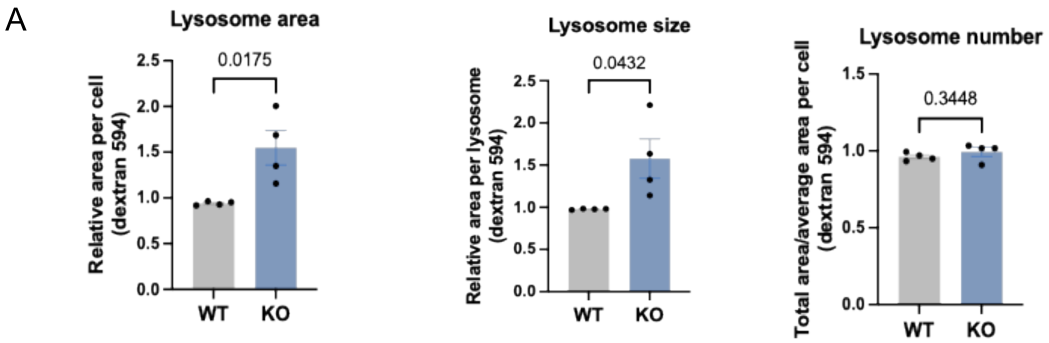


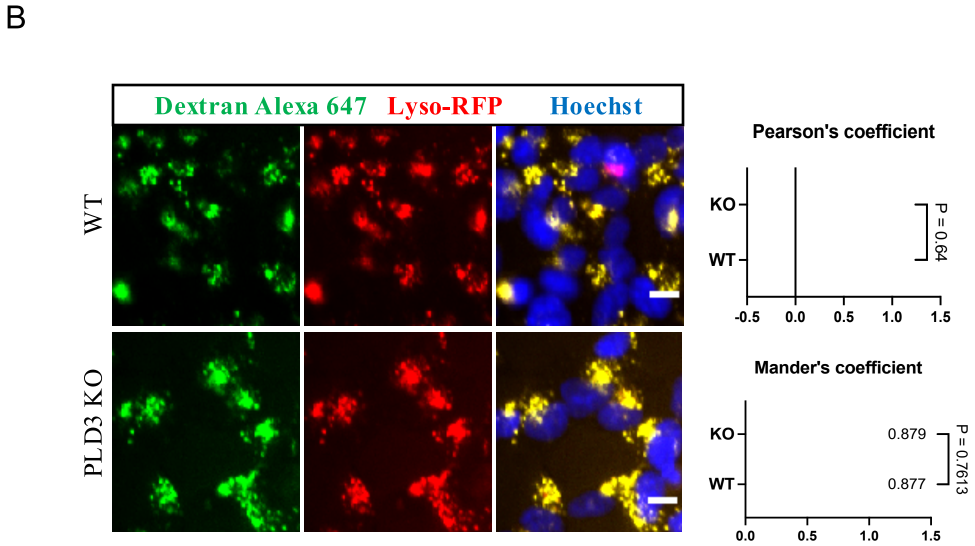

**Fig. S9.** Romero-Fernandez W. and Wang Y. *et al.*, 2025.

A

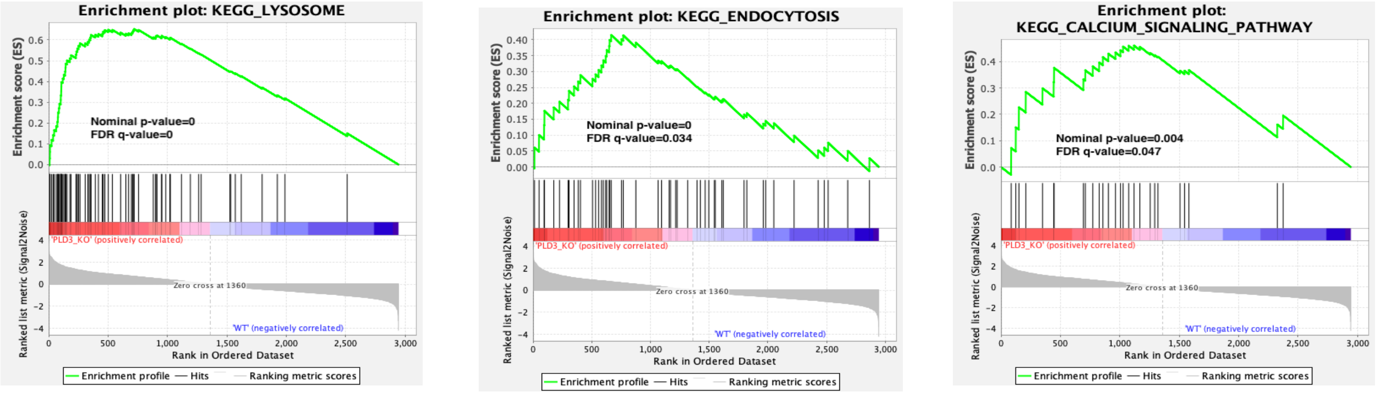


B

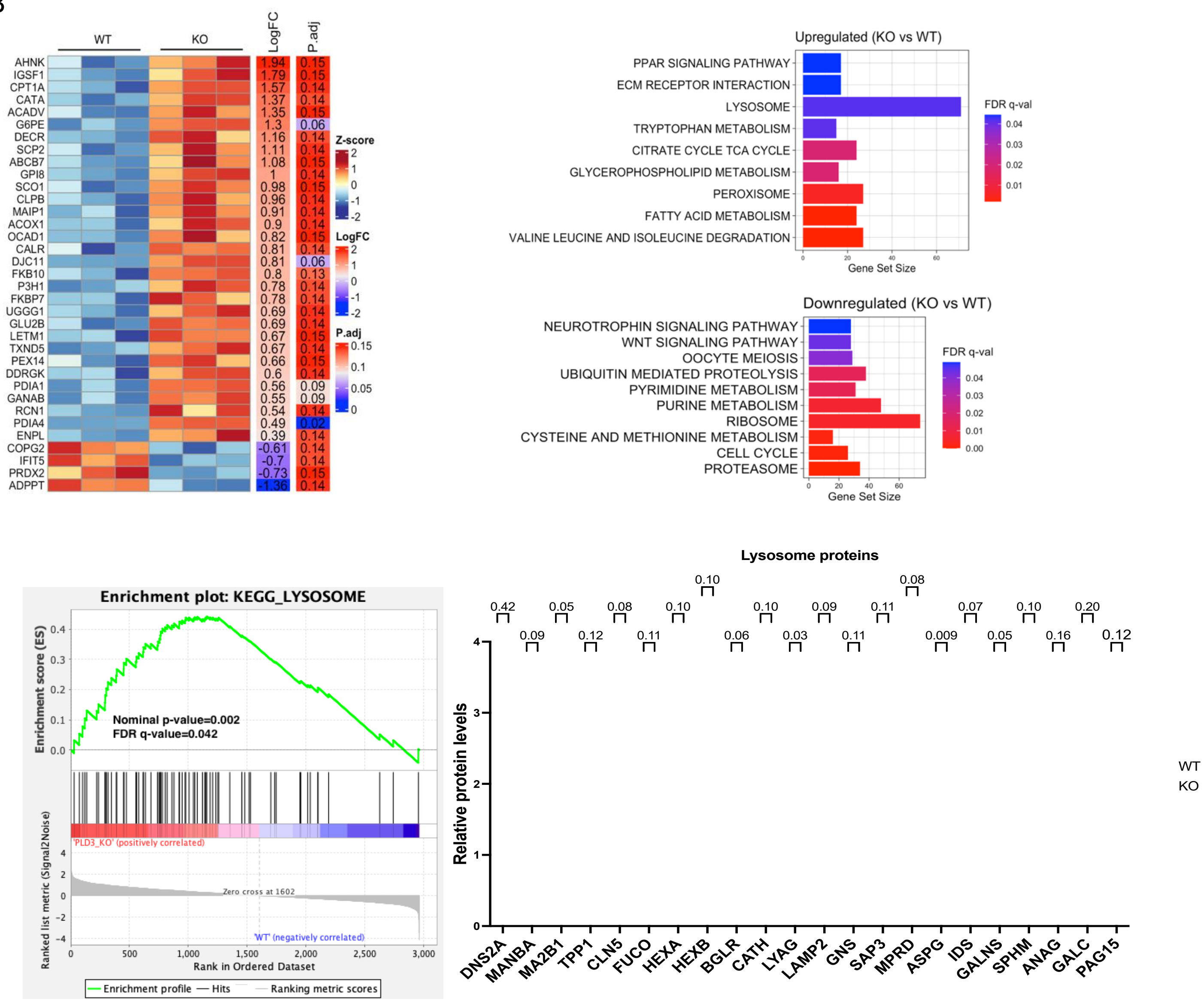

**Fig. S10.** Romero-Fernandez W. and Wang Y. *et al.*, 2025.

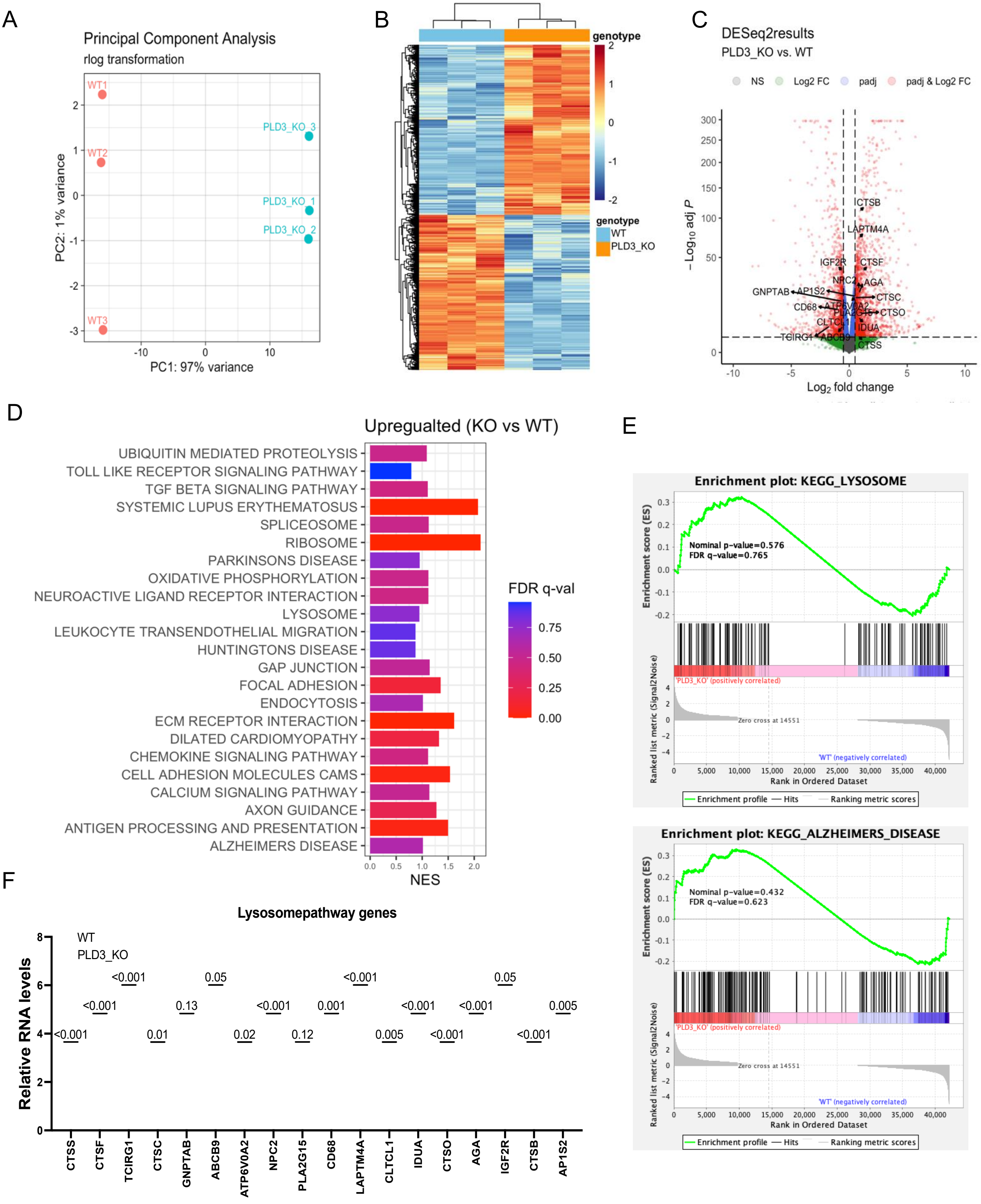

**Fig. S11.** Romero-Fernandez W. and Wang Y. *et al*., 2025.

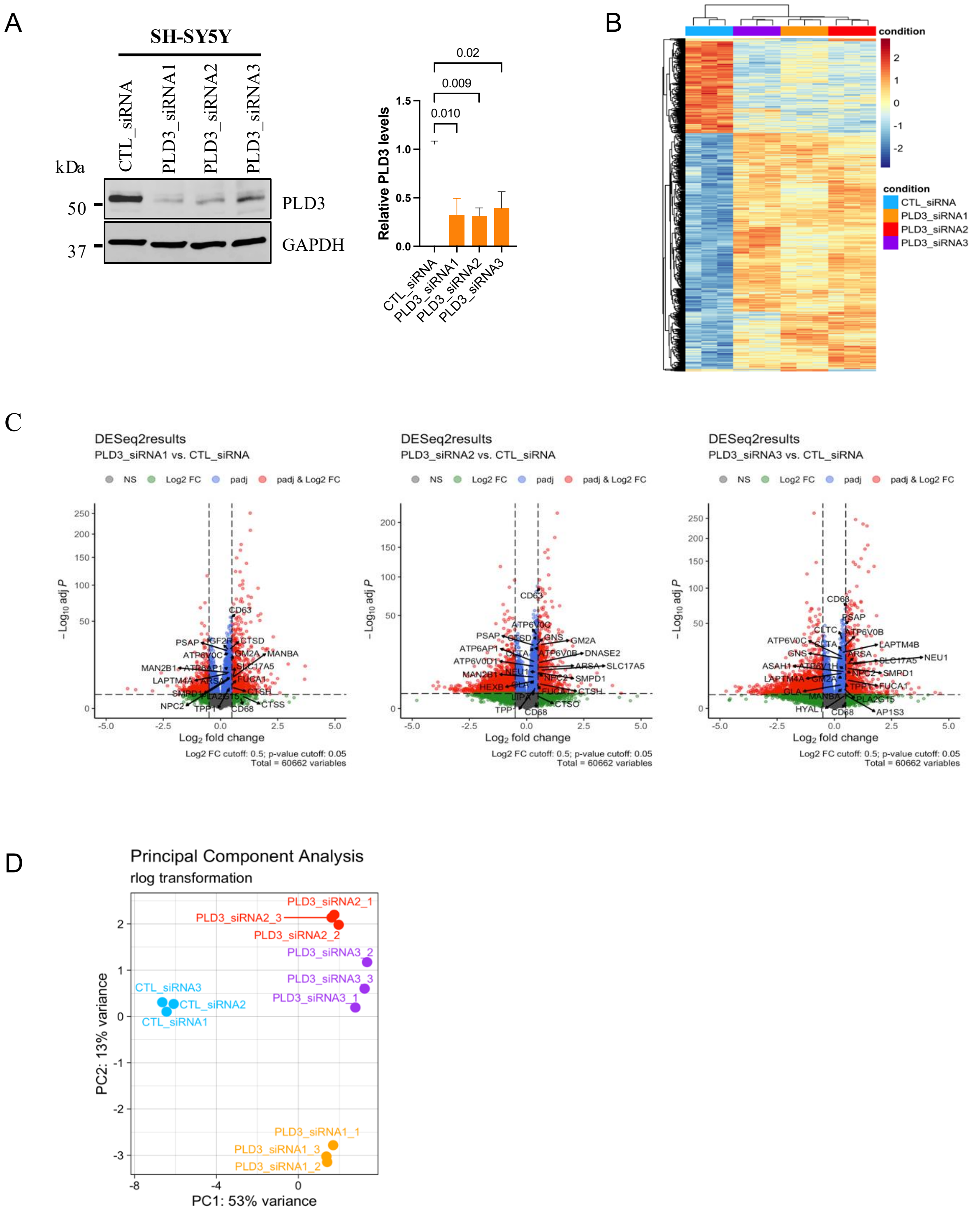

**Fig. S12.** Romero-Fernandez W. and Wang Y. *et al*., 2025.

A

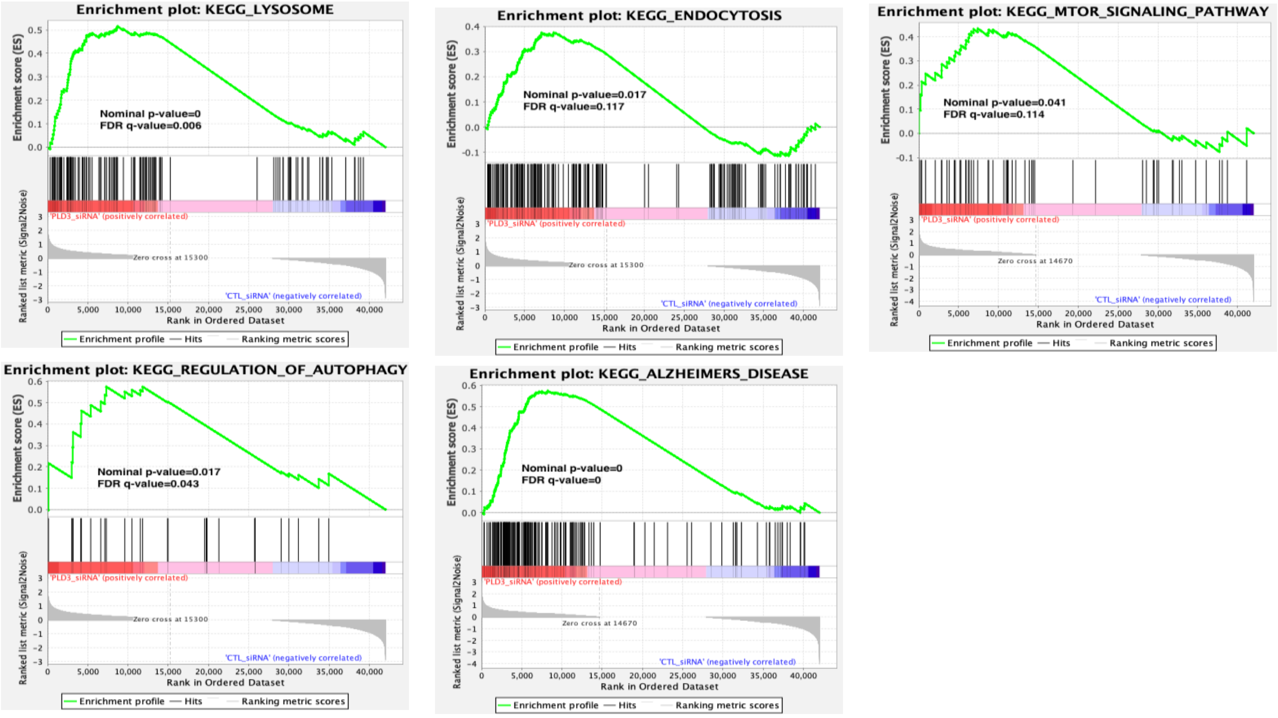

**Fig. S13.** Romero-Fernandez W. and Wang Y. *et al.*, 2025.

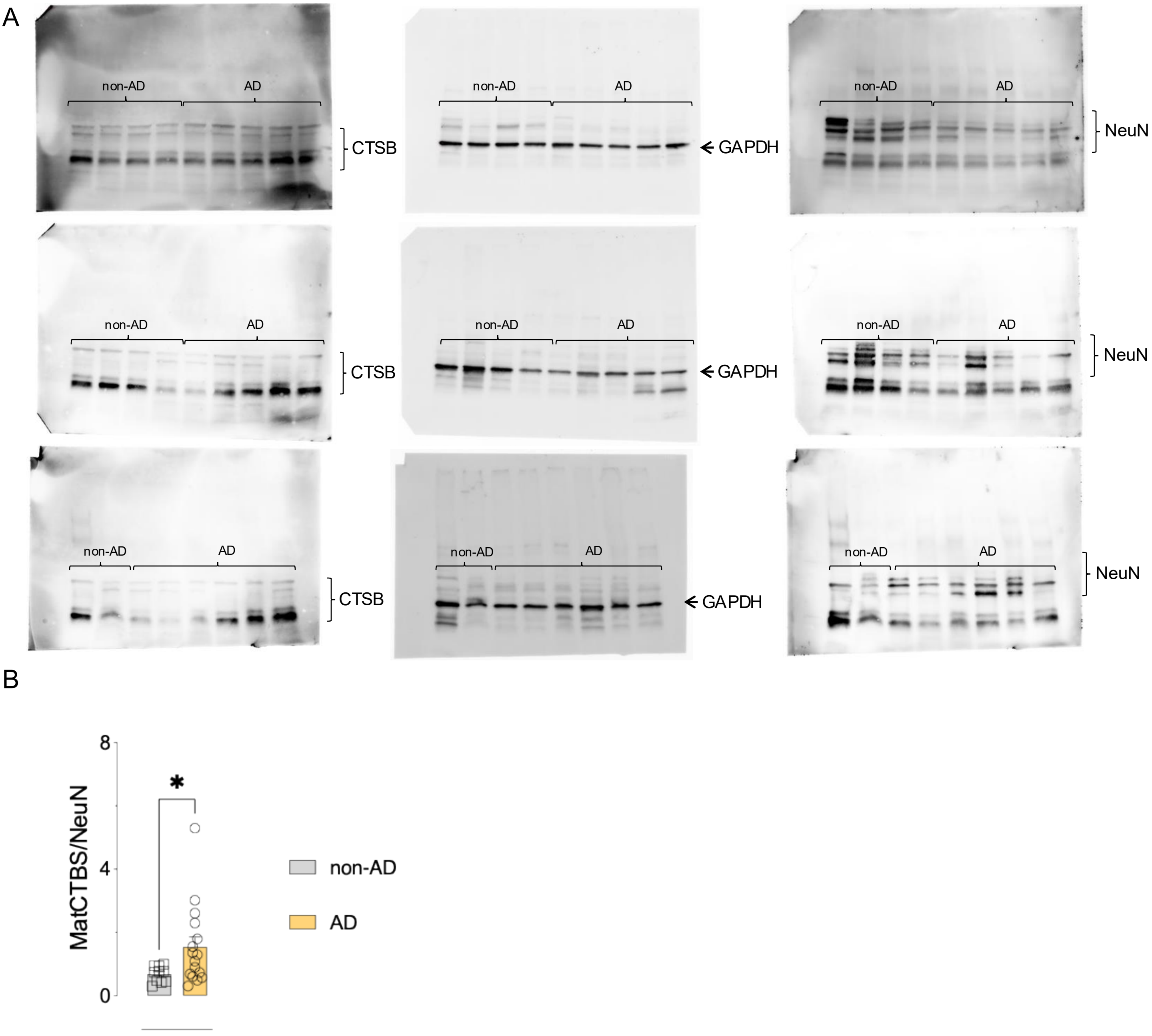

**Fig. S14.** Romero-Fernandez W. and Wang Y. *et al.*, 2025.

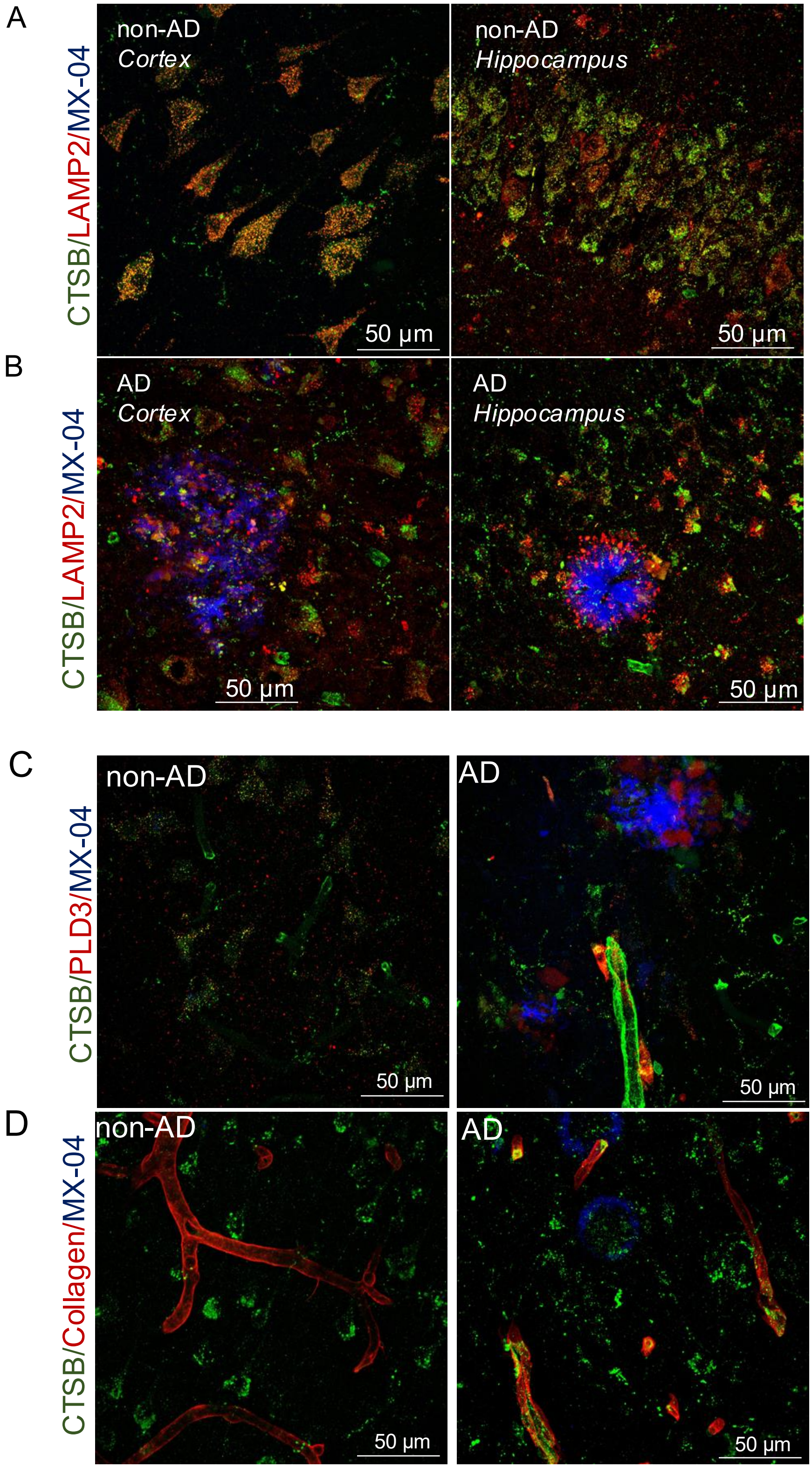

**Fig. S15.** Romero-Fernandez W. and Wang Y. *et al.*, 2025.

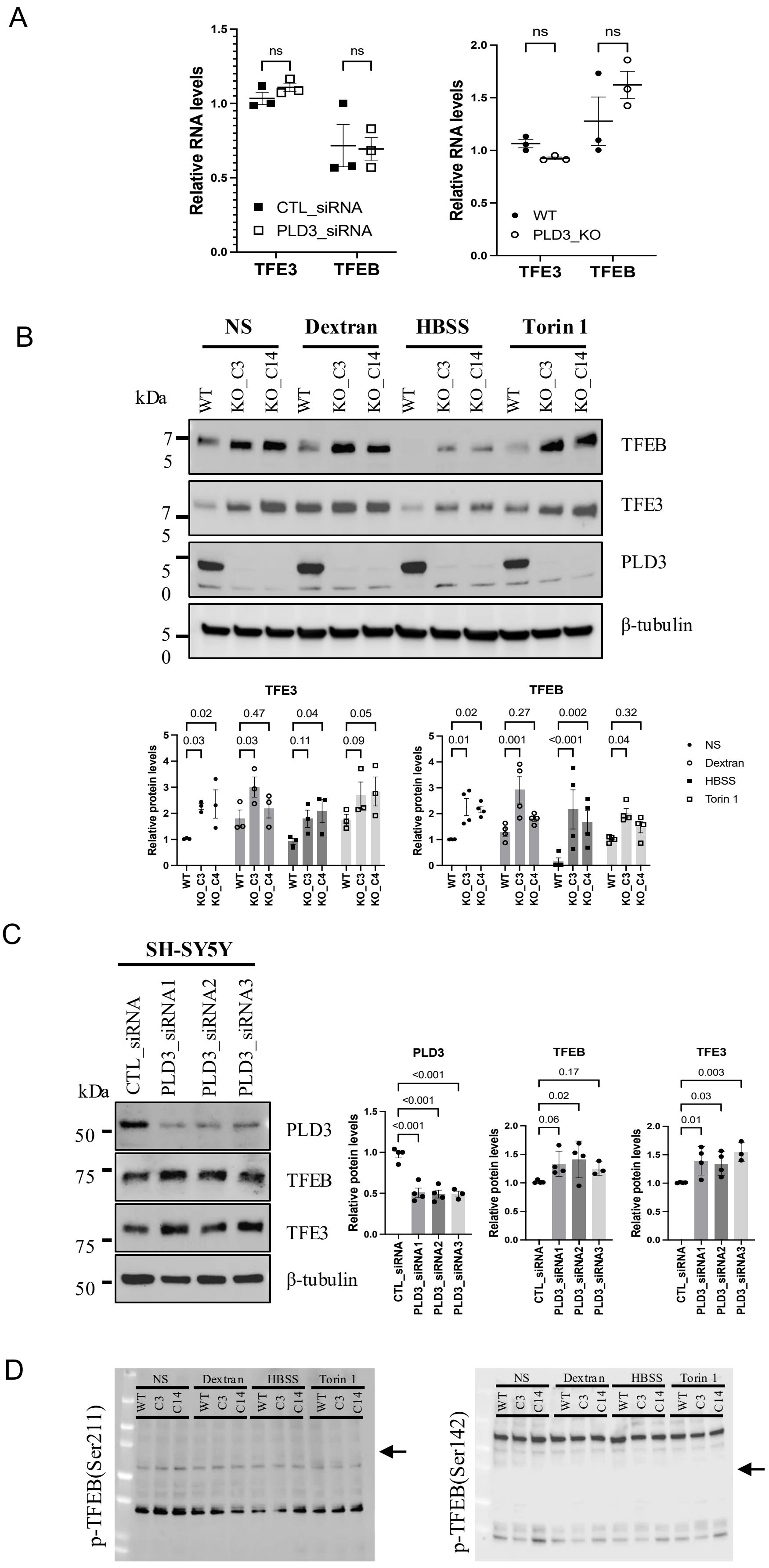

**Fig. S16.** Romero-Fernandez W. and Wang Y. *et al.*, 2025.

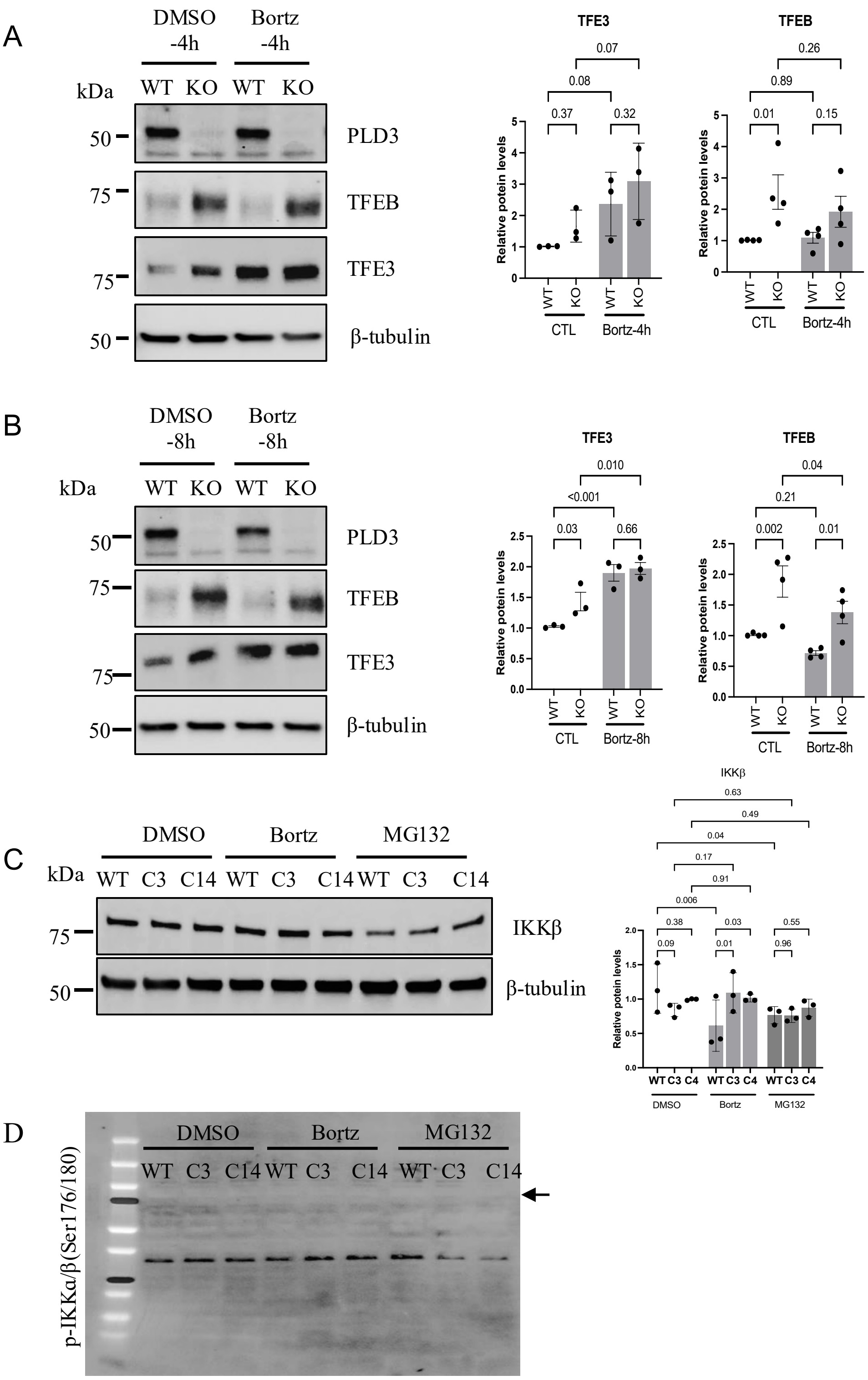

**Fig. S17.** Romero-Fernandez W. and Wang Y. *et al.*, 2025.

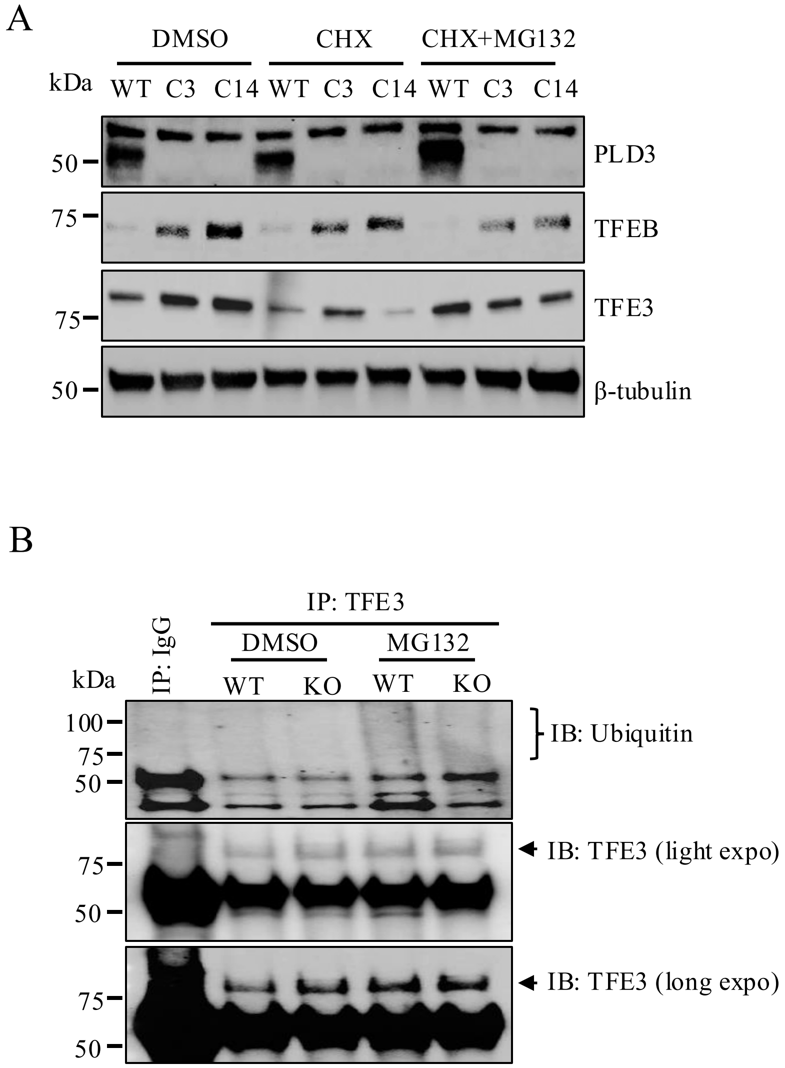

**Fig S18.** Romero-Fernandez W. and Wang Y. *et al.*, 2025.

A

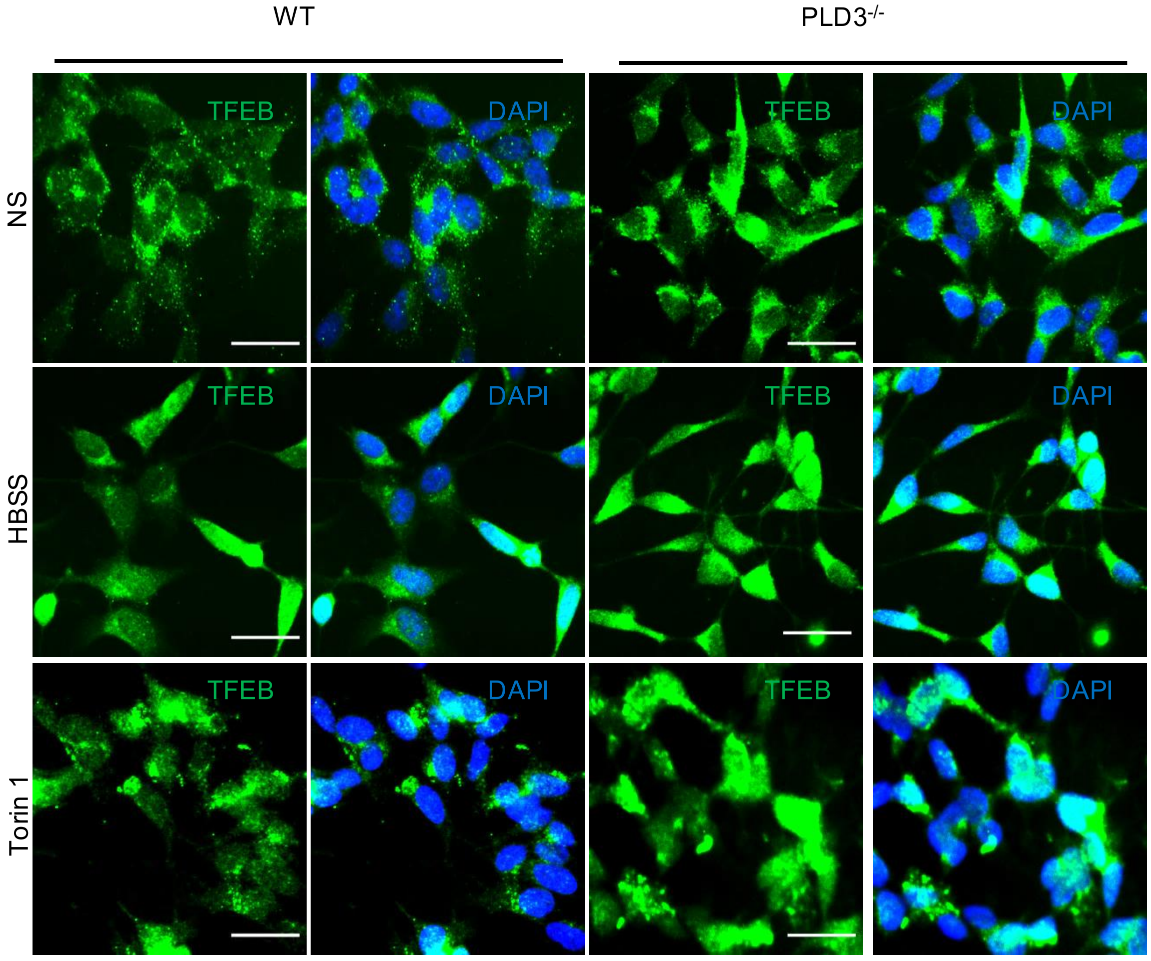
